## Supplementary Table 1 for "Tissue- and population-level microbiome analysis of the wasp spider *Argiope bruennichi* identifies a novel dominant bacterial symbiont"

**Additional File 1: Taxonomy Table**

A table containing the taxonomy of the 573 most abundant amplicon sequence variants obtained by sequencing the microbiome of *Argiope bruennichi*

| **Kingdom** | **Phylum** | **Class** | **Order** | **Family** | **Genus** |
| --- | --- | --- | --- | --- | --- |
| Bacteria | Actinobacteria | Actinobacteria | Micrococcales | Micrococcaceae | Micrococcus |
| Bacteria | Actinobacteria | Actinobacteria | Micrococcales | Microbacteriaceae | Frigoribacterium |
| Bacteria | Actinobacteria | Actinobacteria | Micrococcales | Microbacteriaceae | Curtobacterium |
| Bacteria | Actinobacteria | Actinobacteria | Corynebacteriales | Tsukamurellaceae | Tsukamurella |
| Bacteria | Actinobacteria | Actinobacteria | Micrococcales | Microbacteriaceae | Rathayibacter |
| Bacteria | Actinobacteria | Actinobacteria | Corynebacteriales | Nocardiaceae | Rhodococcus |
| Bacteria | Actinobacteria | Actinobacteria | Streptomycetales | Streptomycetaceae | Streptomyces |
| Bacteria | Actinobacteria | Actinobacteria | Micrococcales | Promicromonosporaceae | Cellulosimicrobium |
| Bacteria | Actinobacteria | Actinobacteria | Frankiales | Nakamurellaceae | Nakamurella |
| Bacteria | Actinobacteria | Actinobacteria | Corynebacteriales | Nocardiaceae | Rhodococcus |
| Bacteria | Actinobacteria | Actinobacteria | Frankiales | Geodermatophilaceae | Geodermatophilus |
| Bacteria | Actinobacteria | Actinobacteria | Micrococcales | Microbacteriaceae | Frigoribacterium |
| Bacteria | Actinobacteria | Actinobacteria | Corynebacteriales | Mycobacteriaceae | Mycobacterium |
| Bacteria | Actinobacteria | Actinobacteria | Corynebacteriales | Corynebacteriaceae | Corynebacterium_1 |
| Bacteria | Actinobacteria | Actinobacteria | Micrococcales | Microbacteriaceae | Leifsonia |
| Bacteria | Actinobacteria | Actinobacteria | Micrococcales | Microbacteriaceae | Amnibacterium |
| Bacteria | Actinobacteria | Actinobacteria | Corynebacteriales | Nocardiaceae | Rhodococcus |
| Bacteria | Actinobacteria | Actinobacteria | Micrococcales | Microbacteriaceae | Amnibacterium |
| Bacteria | Actinobacteria | Actinobacteria | Corynebacteriales | Nocardiaceae | Williamsia |
| Bacteria | Actinobacteria | Actinobacteria | Corynebacteriales | Mycobacteriaceae | Mycobacterium |
| Bacteria | Actinobacteria | Actinobacteria | Corynebacteriales | Corynebacteriaceae | Corynebacterium_1 |
| Bacteria | Actinobacteria | Actinobacteria | Micrococcales | Microbacteriaceae | Amnibacterium |
| Bacteria | Actinobacteria | Actinobacteria | Micrococcales | Brevibacteriaceae | Brevibacterium |
| Bacteria | Actinobacteria | Actinobacteria | Streptomycetales | Streptomycetaceae | Streptomyces |
| Bacteria | Actinobacteria | Actinobacteria | Micrococcales | Microbacteriaceae | NA |
| Bacteria | Actinobacteria | Actinobacteria | Micrococcales | Intrasporangiaceae | NA |
| Bacteria | Actinobacteria | Actinobacteria | Corynebacteriales | Corynebacteriaceae | Corynebacterium |
| Bacteria | Actinobacteria | Actinobacteria | Micrococcales | Microbacteriaceae | Clavibacter |
| Bacteria | Actinobacteria | Actinobacteria | Micrococcales | Microbacteriaceae | Leifsonia |
| Bacteria | Actinobacteria | Actinobacteria | Micrococcales | Intrasporangiaceae | Intrasporangium |
| Bacteria | Actinobacteria | Actinobacteria | Micrococcales | Microbacteriaceae | Agromyces |
| Bacteria | Actinobacteria | Actinobacteria | Frankiales | Nakamurellaceae | Nakamurella |
| Bacteria | Actinobacteria | Actinobacteria | Kineosporiales | Kineosporiaceae | Kineococcus |
| Bacteria | Actinobacteria | Actinobacteria | Propionibacteriales | Nocardioidaceae | Nocardioides |
| Bacteria | Actinobacteria | Actinobacteria | Corynebacteriales | Corynebacteriaceae | Corynebacterium_1 |
| Bacteria | Actinobacteria | Actinobacteria | Micrococcales | Brevibacteriaceae | Brevibacterium |
| Bacteria | Actinobacteria | Actinobacteria | Corynebacteriales | Corynebacteriaceae | Corynebacterium_1 |
| Bacteria | Actinobacteria | Actinobacteria | Pseudonocardiales | Pseudonocardiaceae | Pseudonocardia |
| Bacteria | Actinobacteria | Actinobacteria | Micrococcales | Intrasporangiaceae | Ornithinimicrobium |
| Bacteria | Actinobacteria | Actinobacteria | Corynebacteriales | Corynebacteriaceae | Corynebacterium_1 |
| Bacteria | Actinobacteria | Actinobacteria | Propionibacteriales | Nocardioidaceae | Marmoricola |
| Bacteria | Actinobacteria | Actinobacteria | Micromonosporales | Micromonosporaceae | NA |
| Bacteria | Actinobacteria | Actinobacteria | Pseudonocardiales | Pseudonocardiaceae | Amycolatopsis |
| Bacteria | Actinobacteria | Actinobacteria | Corynebacteriales | Mycobacteriaceae | Mycobacterium |
| Bacteria | Actinobacteria | Actinobacteria | Actinomycetales | Actinomycetaceae | Actinomyces |
| Bacteria | Actinobacteria | Actinobacteria | Propionibacteriales | Nocardioidaceae | Nocardioides |
| Bacteria | Actinobacteria | Actinobacteria | Kineosporiales | Kineosporiaceae | Kineococcus |
| Bacteria | Actinobacteria | Actinobacteria | Frankiales | Frankiaceae | Jatrophihabitans |
| Bacteria | Actinobacteria | Actinobacteria | Corynebacteriales | Corynebacteriaceae | Corynebacterium |
| Bacteria | Actinobacteria | Actinobacteria | Propionibacteriales | Nocardioidaceae | Marmoricola |
| Bacteria | Actinobacteria | Actinobacteria | Micrococcales | Beutenbergiaceae | Salana |
| Bacteria | Actinobacteria | Actinobacteria | Micrococcales | Micrococcaceae | Kocuria |
| Bacteria | Actinobacteria | Actinobacteria | Corynebacteriales | Corynebacteriaceae | Lawsonella |
| Bacteria | Actinobacteria | Actinobacteria | Corynebacteriales | Corynebacteriaceae | Corynebacterium_1 |
| Bacteria | Actinobacteria | Actinobacteria | Propionibacteriales | Nocardioidaceae | Nocardioides |
| Bacteria | Actinobacteria | Actinobacteria | Micrococcales | Micrococcaceae | Pseudarthrobacter |
| Bacteria | Actinobacteria | Actinobacteria | Micrococcales | Microbacteriaceae | Leucobacter |
| Bacteria | Actinobacteria | Actinobacteria | Frankiales | Frankiaceae | Jatrophihabitans |
| Bacteria | Actinobacteria | Actinobacteria | Propionibacteriales | Nocardioidaceae | Nocardioides |
| Bacteria | Actinobacteria | Actinobacteria | Micrococcales | Intrasporangiaceae | Janibacter |
| Bacteria | Actinobacteria | Actinobacteria | Corynebacteriales | Corynebacteriaceae | Corynebacterium_1 |
| Bacteria | Actinobacteria | Actinobacteria | Corynebacteriales | Corynebacteriaceae | Corynebacterium_1 |
| Bacteria | Actinobacteria | Actinobacteria | Corynebacteriales | Nocardiaceae | Rhodococcus |
| Bacteria | Actinobacteria | Actinobacteria | Bifidobacteriales | Bifidobacteriaceae | Gardnerella |
| Bacteria | Actinobacteria | Actinobacteria | Micrococcales | Microbacteriaceae | Herbiconiux |
| Bacteria | Actinobacteria | Actinobacteria | Micrococcales | Microbacteriaceae | NA |
| Bacteria | Actinobacteria | Actinobacteria | NA | NA | NA |
| Bacteria | Actinobacteria | Actinobacteria | Corynebacteriales | Corynebacteriaceae | Corynebacterium_1 |
| Bacteria | Actinobacteria | Actinobacteria | Micrococcales | Cellulomonadaceae | Cellulomonas |
| Bacteria | Actinobacteria | Actinobacteria | Actinomycetales | Actinomycetaceae | Actinomyces |
| Bacteria | Actinobacteria | Actinobacteria | Actinomycetales | Actinomycetaceae | Varibaculum |
| Bacteria | Actinobacteria | Actinobacteria | Micrococcales | Microbacteriaceae | NA |
| Bacteria | Actinobacteria | Actinobacteria | Actinomycetales | Actinomycetaceae | Actinomyces |
| Bacteria | Actinobacteria | Actinobacteria | Micrococcales | Micrococcaceae | Rothia |
| Bacteria | Actinobacteria | Actinobacteria | Micrococcales | Cellulomonadaceae | Oerskovia |
| Bacteria | Proteobacteria | Alphaproteobacteria | Sphingomonadales | Sphingomonadaceae | Sphingomonas |
| Bacteria | Proteobacteria | Alphaproteobacteria | Rhizobiales | Rhizobiaceae | Ochrobactrum |
| Bacteria | Proteobacteria | Alphaproteobacteria | Rhizobiales | Rhizobiaceae | Mesorhizobium |
| Bacteria | Proteobacteria | Alphaproteobacteria | Rhizobiales | Beijerinckiaceae | Methylobacterium |
| Bacteria | Proteobacteria | Alphaproteobacteria | Rhizobiales | Beijerinckiaceae | Bosea |
| Bacteria | Proteobacteria | Alphaproteobacteria | Rhodobacterales | Rhodobacteraceae | NA |
| Bacteria | Proteobacteria | Alphaproteobacteria | Rhizobiales | Beijerinckiaceae | Methylobacterium |
| Bacteria | Proteobacteria | Alphaproteobacteria | Rhizobiales | Beijerinckiaceae | Methylobacterium |
| Bacteria | Proteobacteria | Alphaproteobacteria | Rhodobacterales | Rhodobacteraceae | Paracoccus |
| Bacteria | Proteobacteria | Alphaproteobacteria | Reyranellales | Reyranellaceae | Reyranella |
| Bacteria | Proteobacteria | Alphaproteobacteria | Rhizobiales | Beijerinckiaceae | Methylobacterium |
| Bacteria | Proteobacteria | Alphaproteobacteria | Rhizobiales | Xanthobacteraceae | NA |
| Bacteria | Proteobacteria | Alphaproteobacteria | Rhizobiales | Rhizobiaceae | Ochrobactrum |
| Bacteria | Proteobacteria | Alphaproteobacteria | Rhizobiales | Beijerinckiaceae | 1174-901-12 |
| Bacteria | Proteobacteria | Alphaproteobacteria | Caulobacterales | Caulobacteraceae | Brevundimonas |
| Bacteria | Proteobacteria | Alphaproteobacteria | Sphingomonadales | Sphingomonadaceae | Novosphingobium |
| Bacteria | Proteobacteria | Alphaproteobacteria | Rhizobiales | Rhizobiaceae | Ochrobactrum |
| Bacteria | Proteobacteria | Alphaproteobacteria | Sphingomonadales | Sphingomonadaceae | Stakelama |
| Bacteria | Proteobacteria | Alphaproteobacteria | Rhodobacterales | Rhodobacteraceae | Paracoccus |
| Bacteria | Proteobacteria | Alphaproteobacteria | Caulobacterales | Caulobacteraceae | Caulobacter |
| Bacteria | Proteobacteria | Alphaproteobacteria | Rhizobiales | Beijerinckiaceae | Methylobacterium |
| Bacteria | Proteobacteria | Alphaproteobacteria | Rickettsiales | Mitochondria | NA |
| Bacteria | Proteobacteria | Alphaproteobacteria | Rhizobiales | Beijerinckiaceae | 1174-901-12 |
| Bacteria | Proteobacteria | Alphaproteobacteria | Sphingomonadales | Sphingomonadaceae | Sphingomonas |
| Bacteria | Proteobacteria | Alphaproteobacteria | Sphingomonadales | Sphingomonadaceae | Sphingomonas |
| Bacteria | Proteobacteria | Alphaproteobacteria | Rhizobiales | Xanthobacteraceae | NA |
| Bacteria | Proteobacteria | Alphaproteobacteria | Rhizobiales | Rhizobiaceae | Ochrobactrum |
| Bacteria | Proteobacteria | Alphaproteobacteria | Holosporales | Holosporaceae | NA |
| Bacteria | Proteobacteria | Alphaproteobacteria | Rhizobiales | Rhizobiaceae | Neorhizobium |
| Bacteria | Proteobacteria | Alphaproteobacteria | Rickettsiales | Mitochondria | NA |
| Bacteria | Proteobacteria | Alphaproteobacteria | Rhizobiales | Devosiaceae | Devosia |
| Bacteria | Proteobacteria | Alphaproteobacteria | Sphingomonadales | Sphingomonadaceae | Sphingomonas |
| Bacteria | Proteobacteria | Alphaproteobacteria | Sphingomonadales | Sphingomonadaceae | Ellin6055 |
| Bacteria | Proteobacteria | Alphaproteobacteria | Rhizobiales | Devosiaceae | NA |
| Bacteria | Proteobacteria | Alphaproteobacteria | Rhizobiales | Rhizobiaceae | Brucella |
| Bacteria | Proteobacteria | Alphaproteobacteria | Rhizobiales | Rhizobiaceae | Aquamicrobium |
| Bacteria | Proteobacteria | Alphaproteobacteria | Sphingomonadales | Sphingomonadaceae | Sphingomonas |
| Bacteria | Proteobacteria | Alphaproteobacteria | Sphingomonadales | Sphingomonadaceae | Sphingomonas |
| Bacteria | Proteobacteria | Alphaproteobacteria | Sphingomonadales | Sphingomonadaceae | Sphingobium |
| Bacteria | Proteobacteria | Alphaproteobacteria | Rhizobiales | Rhizobiaceae | Ochrobactrum |
| Bacteria | Proteobacteria | Alphaproteobacteria | Rhizobiales | Beijerinckiaceae | Methylobacterium |
| Bacteria | Proteobacteria | Alphaproteobacteria | Rickettsiales | Mitochondria | NA |
| Bacteria | Proteobacteria | Alphaproteobacteria | Sphingomonadales | Sphingomonadaceae | Sphingomonas |
| Bacteria | Proteobacteria | Alphaproteobacteria | Rhodobacterales | Rhodobacteraceae | Paracoccus |
| Bacteria | Proteobacteria | Alphaproteobacteria | Rhodobacterales | Rhodobacteraceae | Paracoccus |
| Bacteria | Proteobacteria | Alphaproteobacteria | Rhizobiales | Rhizobiaceae | Brucella |
| Bacteria | Proteobacteria | Alphaproteobacteria | Rhizobiales | Beijerinckiaceae | Microvirga |
| Bacteria | Proteobacteria | Alphaproteobacteria | Rhizobiales | Rhizobiaceae | NA |
| Bacteria | Proteobacteria | Alphaproteobacteria | Reyranellales | Reyranellaceae | Reyranella |
| Bacteria | Proteobacteria | Alphaproteobacteria | Rickettsiales | Mitochondria | NA |
| Bacteria | Proteobacteria | Alphaproteobacteria | Rhizobiales | Hyphomicrobiaceae | Pedomicrobium |
| Bacteria | Proteobacteria | Alphaproteobacteria | Sphingomonadales | Sphingomonadaceae | Sphingomonas |
| Bacteria | Proteobacteria | Alphaproteobacteria | Rhizobiales | Beijerinckiaceae | Methylobacterium |
| Bacteria | Proteobacteria | Alphaproteobacteria | Rhizobiales | Rhizobiaceae | Aliihoeflea |
| Bacteria | Proteobacteria | Alphaproteobacteria | Rhizobiales | Xanthobacteraceae | NA |
| Bacteria | Proteobacteria | Alphaproteobacteria | Rickettsiales | Mitochondria | NA |
| Bacteria | Proteobacteria | Alphaproteobacteria | Rickettsiales | Mitochondria | NA |
| Bacteria | Proteobacteria | Alphaproteobacteria | Rhizobiales | Beijerinckiaceae | Methylobacterium |
| Bacteria | Proteobacteria | Alphaproteobacteria | Acetobacterales | Acetobacteraceae | Endobacter |
| Bacteria | Proteobacteria | Alphaproteobacteria | Rhizobiales | Rhizobiaceae | Allorhizobium-Neorhizobium-Pararhizobium-Rhizobium |
| Bacteria | Proteobacteria | Alphaproteobacteria | Rhizobiales | Beijerinckiaceae | Bosea |
| Bacteria | Proteobacteria | Alphaproteobacteria | Rhizobiales | Xanthobacteraceae | NA |
| Bacteria | Proteobacteria | Alphaproteobacteria | Rhizobiales | Beijerinckiaceae | Methylobacterium |
| Bacteria | Proteobacteria | Alphaproteobacteria | Caulobacterales | Caulobacteraceae | Brevundimonas |
| Bacteria | Proteobacteria | Alphaproteobacteria | Rhizobiales | Rhizobiaceae | Aminobacter |
| Bacteria | Proteobacteria | Alphaproteobacteria | Sphingomonadales | Sphingomonadaceae | Sphingomonas |
| Bacteria | Proteobacteria | Alphaproteobacteria | Rhizobiales | Beijerinckiaceae | Qingshengfania |
| Bacteria | Proteobacteria | Alphaproteobacteria | Sphingomonadales | Sphingomonadaceae | Sphingomonas |
| Bacteria | Proteobacteria | Alphaproteobacteria | Rhizobiales | Beijerinckiaceae | Methylobacterium |
| Bacteria | Proteobacteria | Alphaproteobacteria | Rhizobiales | Beijerinckiaceae | Methylobacterium |
| Bacteria | Proteobacteria | Alphaproteobacteria | Rhizobiales | Beijerinckiaceae | Methylobacterium |
| Bacteria | Proteobacteria | Alphaproteobacteria | Acetobacterales | Acetobacteraceae | Roseomonas |
| Bacteria | Proteobacteria | Alphaproteobacteria | Rhizobiales | Rhizobiaceae | Ochrobactrum |
| Bacteria | Proteobacteria | Alphaproteobacteria | Rhizobiales | Beijerinckiaceae | Microvirga |
| Bacteria | Proteobacteria | Alphaproteobacteria | Rhizobiales | Rhizobiaceae | Pseudochrobactrum |
| Bacteria | Proteobacteria | Alphaproteobacteria | Sphingomonadales | Sphingomonadaceae | Sphingomonas |
| Bacteria | Proteobacteria | Alphaproteobacteria | Rickettsiales | Mitochondria | NA |
| Bacteria | Proteobacteria | Alphaproteobacteria | Caulobacterales | Caulobacteraceae | Brevundimonas |
| Bacteria | Proteobacteria | Alphaproteobacteria | Sphingomonadales | Sphingomonadaceae | Sphingorhabdus |
| Bacteria | Proteobacteria | Alphaproteobacteria | Rhizobiales | Rhizobiaceae | Ochrobactrum |
| Bacteria | Proteobacteria | Alphaproteobacteria | Caulobacterales | Caulobacteraceae | Brevundimonas |
| Bacteria | Proteobacteria | Alphaproteobacteria | Rhizobiales | Rhizobiales_Incertae_Sedis | Alsobacter |
| Bacteria | Proteobacteria | Alphaproteobacteria | Sphingomonadales | Sphingomonadaceae | Sphingobium |
| Bacteria | Proteobacteria | Alphaproteobacteria | Rhizobiales | Beijerinckiaceae | Methylobacterium |
| Bacteria | Proteobacteria | Alphaproteobacteria | Rhizobiales | Rhizobiaceae | Ochrobactrum |
| Bacteria | Proteobacteria | Alphaproteobacteria | Holosporales | Holosporaceae | NA |
| Bacteria | Proteobacteria | Alphaproteobacteria | Rhizobiales | Labraceae | Labrys |
| Bacteria | Proteobacteria | Alphaproteobacteria | Rhizobiales | Labraceae | Labrys |
| Bacteria | Proteobacteria | Alphaproteobacteria | Rhizobiales | Labraceae | Labrys |
| Bacteria | Proteobacteria | Alphaproteobacteria | Rhodobacterales | Rhodobacteraceae | Rubellimicrobium |
| Bacteria | Proteobacteria | Alphaproteobacteria | Azospirillales | Azospirillaceae | Skermanella |
| Bacteria | Proteobacteria | Alphaproteobacteria | Sphingomonadales | Sphingomonadaceae | Sphingomonas |
| Bacteria | Proteobacteria | Alphaproteobacteria | Rhodobacterales | Rhodobacteraceae | Amaricoccus |
| Bacteria | Proteobacteria | Alphaproteobacteria | Sphingomonadales | Sphingomonadaceae | Altererythrobacter |
| Bacteria | Proteobacteria | Alphaproteobacteria | Rhizobiales | Devosiaceae | Devosia |
| Bacteria | Proteobacteria | Alphaproteobacteria | Rickettsiales | Mitochondria | NA |
| Bacteria | Firmicutes | Bacilli | Bacillales | Bacillaceae | Bacillus |
| Bacteria | Firmicutes | Bacilli | Lactobacillales | Enterococcaceae | Enterococcus |
| Bacteria | Firmicutes | Bacilli | Bacillales | Planococcaceae | Lysinibacillus |
| Bacteria | Firmicutes | Bacilli | Bacillales | Staphylococcaceae | Staphylococcus |
| Bacteria | Firmicutes | Bacilli | Lactobacillales | Streptococcaceae | Streptococcus |
| Bacteria | Firmicutes | Bacilli | Bacillales | Paenibacillaceae | Paenibacillus |
| Bacteria | Firmicutes | Bacilli | Bacillales | Bacillaceae | Fictibacillus |
| Bacteria | Firmicutes | Bacilli | Lactobacillales | Enterococcaceae | Enterococcus |
| Bacteria | Firmicutes | Bacilli | Lactobacillales | Streptococcaceae | Streptococcus |
| Bacteria | Firmicutes | Bacilli | Bacillales | Planococcaceae | Solibacillus |
| Bacteria | Firmicutes | Bacilli | Lactobacillales | Streptococcaceae | Streptococcus |
| Bacteria | Firmicutes | Bacilli | Bacillales | Paenibacillaceae | Paenibacillus |
| Bacteria | Firmicutes | Bacilli | Bacillales | Staphylococcaceae | Staphylococcus |
| Bacteria | Firmicutes | Bacilli | Bacillales | Family_XI | Gemella |
| Bacteria | Firmicutes | Bacilli | Bacillales | Planococcaceae | Lysinibacillus |
| Bacteria | Firmicutes | Bacilli | Lactobacillales | Enterococcaceae | Enterococcus |
| Bacteria | Firmicutes | Bacilli | Lactobacillales | Enterococcaceae | Enterococcus |
| Bacteria | Firmicutes | Bacilli | Bacillales | Paenibacillaceae | Brevibacillus |
| Bacteria | Firmicutes | Bacilli | Lactobacillales | Enterococcaceae | Enterococcus |
| Bacteria | Firmicutes | Bacilli | Bacillales | Bacillaceae | Bacillus |
| Bacteria | Firmicutes | Bacilli | Bacillales | Bacillaceae | NA |
| Bacteria | Firmicutes | Bacilli | Lactobacillales | Streptococcaceae | Streptococcus |
| Bacteria | Firmicutes | Bacilli | Lactobacillales | Leuconostocaceae | Weissella |
| Bacteria | Firmicutes | Bacilli | Bacillales | Planococcaceae | Solibacillus |
| Bacteria | Firmicutes | Bacilli | Lactobacillales | Streptococcaceae | Lactococcus |
| Bacteria | Firmicutes | Bacilli | Lactobacillales | NA | NA |
| Bacteria | Firmicutes | Bacilli | Bacillales | Paenibacillaceae | Paenibacillus |
| Bacteria | Firmicutes | Bacilli | Bacillales | Paenibacillaceae | Brevibacillus |
| Bacteria | Firmicutes | Bacilli | Lactobacillales | Enterococcaceae | Enterococcus |
| Bacteria | Firmicutes | Bacilli | Bacillales | Staphylococcaceae | Staphylococcus |
| Bacteria | Firmicutes | Bacilli | Bacillales | Bacillaceae | Anoxybacillus |
| Bacteria | Firmicutes | Bacilli | Bacillales | Bacillaceae | Oceanobacillus |
| Bacteria | Firmicutes | Bacilli | Bacillales | Staphylococcaceae | Staphylococcus |
| Bacteria | Firmicutes | Bacilli | Bacillales | Family_XII | Exiguobacterium |
| Bacteria | Firmicutes | Bacilli | Lactobacillales | Carnobacteriaceae | Dolosigranulum |
| Bacteria | Firmicutes | Bacilli | Bacillales | Bacillaceae | NA |
| Bacteria | Firmicutes | Bacilli | Bacillales | Planococcaceae | Lysinibacillus |
| Bacteria | Firmicutes | Bacilli | Bacillales | Planococcaceae | NA |
| Bacteria | Firmicutes | Bacilli | Lactobacillales | NA | NA |
| Bacteria | Firmicutes | Bacilli | Lactobacillales | Streptococcaceae | Lactococcus |
| Bacteria | Firmicutes | Bacilli | Lactobacillales | Streptococcaceae | Lactococcus |
| Bacteria | Firmicutes | Bacilli | Bacillales | Bacillaceae | Bacillus |
| Bacteria | Firmicutes | Bacilli | Lactobacillales | Lactobacillaceae | Lactobacillus |
| Bacteria | Firmicutes | Bacilli | Lactobacillales | Enterococcaceae | Enterococcus |
| Bacteria | Firmicutes | Bacilli | Bacillales | Bacillaceae | NA |
| Bacteria | Firmicutes | Bacilli | Lactobacillales | Enterococcaceae | Enterococcus |
| Bacteria | Firmicutes | Bacilli | Lactobacillales | Enterococcaceae | Enterococcus |
| Bacteria | Firmicutes | Bacilli | Bacillales | Staphylococcaceae | Staphylococcus |
| Bacteria | Firmicutes | Bacilli | Bacillales | Bacillaceae | Bacillus |
| Bacteria | Firmicutes | Bacilli | Bacillales | Bacillaceae | Bacillus |
| Bacteria | Firmicutes | Bacilli | Bacillales | Bacillaceae | Bacillus |
| Bacteria | Firmicutes | Bacilli | Lactobacillales | Aerococcaceae | Aerococcus |
| Bacteria | Firmicutes | Bacilli | Bacillales | Paenibacillaceae | Paenibacillus |
| Bacteria | Firmicutes | Bacilli | Bacillales | Bacillaceae | Gracilibacillus |
| Bacteria | Firmicutes | Bacilli | Bacillales | Paenibacillaceae | Paenibacillus |
| Bacteria | Firmicutes | Bacilli | Lactobacillales | Aerococcaceae | Facklamia |
| Bacteria | Firmicutes | Bacilli | Bacillales | Paenibacillaceae | Paenibacillus |
| Bacteria | Firmicutes | Bacilli | Lactobacillales | Enterococcaceae | Enterococcus |
| Bacteria | Firmicutes | Bacilli | Lactobacillales | Leuconostocaceae | Leuconostoc |
| Bacteria | Firmicutes | Bacilli | Bacillales | Paenibacillaceae | Oxalophagus |
| Bacteria | Bacteroidetes | Bacteroidia | Flavobacteriales | Weeksellaceae | NA |
| Bacteria | Bacteroidetes | Bacteroidia | Sphingobacteriales | Sphingobacteriaceae | Pedobacter |
| Bacteria | Bacteroidetes | Bacteroidia | Flavobacteriales | Weeksellaceae | NA |
| Bacteria | Bacteroidetes | Bacteroidia | Chitinophagales | Chitinophagaceae | Sediminibacterium |
| Bacteria | Bacteroidetes | Bacteroidia | Sphingobacteriales | Sphingobacteriaceae | Sphingobacterium |
| Bacteria | Bacteroidetes | Bacteroidia | Flavobacteriales | Weeksellaceae | NA |
| Bacteria | Bacteroidetes | Bacteroidia | Flavobacteriales | Weeksellaceae | Chryseobacterium |
| Bacteria | Bacteroidetes | Bacteroidia | Bacteroidales | Paludibacteraceae | Paludibacter |
| Bacteria | Bacteroidetes | Bacteroidia | Flavobacteriales | Weeksellaceae | Chryseobacterium |
| Bacteria | Bacteroidetes | Bacteroidia | Chitinophagales | Chitinophagaceae | Sediminibacterium |
| Bacteria | Bacteroidetes | Bacteroidia | Flavobacteriales | Weeksellaceae | Elizabethkingia |
| Bacteria | Bacteroidetes | Bacteroidia | Flavobacteriales | Weeksellaceae | Chryseobacterium |
| Bacteria | Bacteroidetes | Bacteroidia | Flavobacteriales | Weeksellaceae | Chryseobacterium |
| Bacteria | Bacteroidetes | Bacteroidia | Cytophagales | Spirosomaceae | Dyadobacter |
| Bacteria | Bacteroidetes | Bacteroidia | Cytophagales | Hymenobacteraceae | Hymenobacter |
| Bacteria | Bacteroidetes | Bacteroidia | Flavobacteriales | Weeksellaceae | Chryseobacterium |
| Bacteria | Bacteroidetes | Bacteroidia | Cytophagales | Hymenobacteraceae | Hymenobacter |
| Bacteria | Bacteroidetes | Bacteroidia | Cytophagales | Hymenobacteraceae | Hymenobacter |
| Bacteria | Bacteroidetes | Bacteroidia | Sphingobacteriales | Sphingobacteriaceae | Pedobacter |
| Bacteria | Bacteroidetes | Bacteroidia | Cytophagales | Microscillaceae | NA |
| Bacteria | Bacteroidetes | Bacteroidia | Flavobacteriales | Weeksellaceae | Chryseobacterium |
| Bacteria | Bacteroidetes | Bacteroidia | Flavobacteriales | Weeksellaceae | Bergeyella |
| Bacteria | Bacteroidetes | Bacteroidia | Cytophagales | Hymenobacteraceae | Hymenobacter |
| Bacteria | Bacteroidetes | Bacteroidia | Flavobacteriales | Flavobacteriaceae | Flavobacterium |
| Bacteria | Bacteroidetes | Bacteroidia | Cytophagales | Hymenobacteraceae | Hymenobacter |
| Bacteria | Bacteroidetes | Bacteroidia | Flavobacteriales | Weeksellaceae | Chryseobacterium |
| Bacteria | Bacteroidetes | Bacteroidia | Bacteroidales | Rikenellaceae | NA |
| Bacteria | Bacteroidetes | Bacteroidia | Flavobacteriales | Weeksellaceae | Cloacibacterium |
| Bacteria | Bacteroidetes | Bacteroidia | Flavobacteriales | Weeksellaceae | Chryseobacterium |
| Bacteria | Bacteroidetes | Bacteroidia | Chitinophagales | Saprospiraceae | OLB8 |
| Bacteria | Bacteroidetes | Bacteroidia | Flavobacteriales | Weeksellaceae | Chryseobacterium |
| Bacteria | Bacteroidetes | Bacteroidia | Cytophagales | Spirosomaceae | Spirosoma |
| Bacteria | Bacteroidetes | Bacteroidia | Flavobacteriales | Weeksellaceae | Chryseobacterium |
| Bacteria | Bacteroidetes | Bacteroidia | Sphingobacteriales | Sphingobacteriaceae | Sphingobacterium |
| Bacteria | Bacteroidetes | Bacteroidia | Flavobacteriales | Weeksellaceae | Cloacibacterium |
| Bacteria | Bacteroidetes | Bacteroidia | Chitinophagales | Chitinophagaceae | Sediminibacterium |
| Bacteria | Bacteroidetes | Bacteroidia | Cytophagales | Hymenobacteraceae | Hymenobacter |
| Bacteria | Bacteroidetes | Bacteroidia | Sphingobacteriales | Sphingobacteriaceae | Sphingobacterium |
| Bacteria | Bacteroidetes | Bacteroidia | Flavobacteriales | Flavobacteriaceae | Flavobacterium |
| Bacteria | Bacteroidetes | Bacteroidia | Sphingobacteriales | Sphingobacteriaceae | Sphingobacterium |
| Bacteria | Bacteroidetes | Bacteroidia | Flavobacteriales | Weeksellaceae | Chryseobacterium |
| Bacteria | Bacteroidetes | Bacteroidia | Sphingobacteriales | Sphingobacteriaceae | Pedobacter |
| Bacteria | Bacteroidetes | Bacteroidia | Flavobacteriales | Flavobacteriaceae | Flavobacterium |
| Bacteria | Bacteroidetes | Bacteroidia | Flavobacteriales | Flavobacteriaceae | Wenyingzhuangia |
| Bacteria | Bacteroidetes | Bacteroidia | Bacteroidales | Prevotellaceae | Prevotella |
| Bacteria | Bacteroidetes | Bacteroidia | Bacteroidales | Dysgonomonadaceae | Dysgonomonas |
| Bacteria | Bacteroidetes | Bacteroidia | Cytophagales | Hymenobacteraceae | Hymenobacter |
| Bacteria | Bacteroidetes | Bacteroidia | Sphingobacteriales | Sphingobacteriaceae | Pedobacter |
| Bacteria | Bacteroidetes | Bacteroidia | Chitinophagales | Chitinophagaceae | NA |
| Bacteria | Firmicutes | Clostridia | Clostridiales | Clostridiaceae_1 | Clostridium_sensu_stricto_16 |
| Bacteria | Firmicutes | Clostridia | Clostridiales | Clostridiaceae_1 | Clostridium_sensu_stricto_7 |
| Bacteria | Firmicutes | Clostridia | Clostridiales | Clostridiaceae_1 | Clostridium_sensu_stricto_7 |
| Bacteria | Firmicutes | Clostridia | Clostridiales | Clostridiaceae_1 | Clostridium_sensu_stricto_7 |
| Bacteria | Firmicutes | Clostridia | Clostridiales | Clostridiaceae_1 | Clostridium_sensu_stricto_7 |
| Bacteria | Firmicutes | Clostridia | Clostridiales | Clostridiaceae_1 | Clostridium_sensu_stricto_7 |
| Bacteria | Firmicutes | Clostridia | Clostridiales | NA | NA |
| Bacteria | Firmicutes | Clostridia | Clostridiales | Clostridiaceae_1 | Clostridium_sensu_stricto_7 |
| Bacteria | Firmicutes | Clostridia | Clostridiales | Clostridiaceae_1 | Clostridium_sensu_stricto_7 |
| Bacteria | Firmicutes | Clostridia | Clostridiales | Clostridiaceae_1 | Clostridium_sensu_stricto_7 |
| Bacteria | Firmicutes | Clostridia | Clostridiales | Clostridiaceae_1 | Clostridium_sensu_stricto_7 |
| Bacteria | Firmicutes | Clostridia | Clostridiales | Clostridiaceae_1 | Clostridium_sensu_stricto_7 |
| Bacteria | Firmicutes | Clostridia | Clostridiales | Clostridiaceae_1 | Clostridium_sensu_stricto_7 |
| Bacteria | Firmicutes | Clostridia | Clostridiales | Clostridiaceae_1 | Clostridium_sensu_stricto_7 |
| Bacteria | Firmicutes | Clostridia | Clostridiales | Clostridiaceae_1 | Clostridium_sensu_stricto_7 |
| Bacteria | Firmicutes | Clostridia | Clostridiales | Clostridiaceae_1 | Clostridium_sensu_stricto_7 |
| Bacteria | Firmicutes | Clostridia | Clostridiales | Clostridiaceae_1 | Clostridium_sensu_stricto_7 |
| Bacteria | Firmicutes | Clostridia | Clostridiales | Clostridiaceae_1 | Clostridium_sensu_stricto_16 |
| Bacteria | Firmicutes | Clostridia | Clostridiales | Clostridiaceae_1 | Clostridium_sensu_stricto_7 |
| Bacteria | Firmicutes | Clostridia | Clostridiales | Clostridiaceae_1 | Clostridium_sensu_stricto_7 |
| Bacteria | Firmicutes | Clostridia | Clostridiales | Clostridiaceae_1 | Clostridium_sensu_stricto_7 |
| Bacteria | Firmicutes | Clostridia | Clostridiales | Clostridiaceae_1 | Clostridium_sensu_stricto_7 |
| Bacteria | Firmicutes | Clostridia | Clostridiales | Clostridiaceae_1 | Clostridium_sensu_stricto_7 |
| Bacteria | Firmicutes | Clostridia | Clostridiales | Clostridiaceae_1 | Clostridium_sensu_stricto_7 |
| Bacteria | Firmicutes | Clostridia | Clostridiales | Ruminococcaceae | NA |
| Bacteria | Firmicutes | Clostridia | Clostridiales | Clostridiaceae_1 | Clostridium_sensu_stricto_7 |
| Bacteria | Firmicutes | Clostridia | Clostridiales | Clostridiaceae_1 | Clostridium_sensu_stricto_3 |
| Bacteria | Firmicutes | Clostridia | Clostridiales | Clostridiaceae_1 | Clostridium_sensu_stricto_3 |
| Bacteria | Firmicutes | Clostridia | Clostridiales | Clostridiaceae_1 | Clostridium_sensu_stricto_7 |
| Bacteria | Firmicutes | Clostridia | Clostridiales | Clostridiaceae_1 | Clostridium_sensu_stricto_7 |
| Bacteria | Firmicutes | Clostridia | Clostridiales | Clostridiaceae_1 | Clostridium_sensu_stricto_7 |
| Bacteria | Firmicutes | Clostridia | Clostridiales | Clostridiaceae_1 | Clostridium_sensu_stricto_7 |
| Bacteria | Firmicutes | Clostridia | Clostridiales | Clostridiaceae_1 | Clostridium_sensu_stricto_7 |
| Bacteria | Firmicutes | Clostridia | Clostridiales | Clostridiaceae_1 | Clostridium_sensu_stricto_7 |
| Bacteria | Firmicutes | Clostridia | Clostridiales | Clostridiaceae_1 | Clostridium_sensu_stricto_3 |
| Bacteria | Firmicutes | Clostridia | Clostridiales | Clostridiaceae_1 | Clostridium_sensu_stricto_7 |
| Bacteria | Firmicutes | Clostridia | Clostridiales | Clostridiaceae_1 | Clostridium_sensu_stricto_3 |
| Bacteria | Firmicutes | Clostridia | Clostridiales | Ruminococcaceae | CAG-352 |
| Bacteria | Firmicutes | Clostridia | Clostridiales | Clostridiaceae_1 | Clostridium_sensu_stricto_7 |
| Bacteria | Firmicutes | Clostridia | Clostridiales | Clostridiaceae_1 | Clostridium_sensu_stricto_7 |
| Bacteria | Firmicutes | Clostridia | Clostridiales | Clostridiaceae_1 | Clostridium_sensu_stricto_7 |
| Bacteria | Firmicutes | Clostridia | Clostridiales | Lachnospiraceae | Tyzzerella |
| Bacteria | Firmicutes | Clostridia | Clostridiales | Clostridiaceae_1 | Clostridium_sensu_stricto_7 |
| Bacteria | Firmicutes | Clostridia | Clostridiales | Clostridiaceae_1 | Clostridium_sensu_stricto_7 |
| Bacteria | Firmicutes | Clostridia | Clostridiales | Clostridiaceae_1 | Clostridium_sensu_stricto_7 |
| Bacteria | Firmicutes | Clostridia | Clostridiales | Clostridiaceae_1 | Clostridium_sensu_stricto_8 |
| Bacteria | Firmicutes | Clostridia | Clostridiales | Clostridiaceae_1 | Clostridium_sensu_stricto_7 |
| Bacteria | Firmicutes | Clostridia | Clostridiales | Clostridiaceae_1 | Clostridium_sensu_stricto_7 |
| Bacteria | Firmicutes | Clostridia | Clostridiales | Clostridiaceae_1 | Clostridium_sensu_stricto_7 |
| Bacteria | Firmicutes | Clostridia | Clostridiales | Peptostreptococcaceae | Paraclostridium |
| Bacteria | Firmicutes | Clostridia | Clostridiales | Lachnospiraceae | Anaerosporobacter |
| Bacteria | Firmicutes | Clostridia | Clostridiales | Clostridiaceae_1 | Clostridium_sensu_stricto_3 |
| Bacteria | Firmicutes | Clostridia | Clostridiales | Family_XI | Anaerococcus |
| Bacteria | Firmicutes | Clostridia | Clostridiales | Clostridiaceae_1 | Clostridium_sensu_stricto_7 |
| Bacteria | Firmicutes | Clostridia | Clostridiales | Lachnospiraceae | NA |
| Bacteria | Firmicutes | Clostridia | Clostridiales | Lachnospiraceae | NA |
| Bacteria | Firmicutes | Clostridia | Clostridiales | Ruminococcaceae | CAG-352 |
| Bacteria | Firmicutes | Clostridia | Clostridiales | Clostridiaceae_1 | Clostridium_sensu_stricto_7 |
| Bacteria | Firmicutes | Clostridia | NA | NA | NA |
| Bacteria | Firmicutes | Clostridia | Clostridiales | Ruminococcaceae | Caproiciproducens |
| Bacteria | Firmicutes | Clostridia | Clostridiales | Ruminococcaceae | NA |
| Bacteria | Firmicutes | Clostridia | Clostridiales | Peptostreptococcaceae | Romboutsia |
| Bacteria | Firmicutes | Clostridia | Clostridiales | Clostridiaceae_1 | Clostridium_sensu_stricto_16 |
| Bacteria | Firmicutes | Clostridia | Clostridiales | Clostridiaceae_1 | Clostridium_sensu_stricto_8 |
| Bacteria | Firmicutes | Clostridia | Clostridiales | NA | NA |
| Bacteria | Firmicutes | Clostridia | Clostridiales | Clostridiaceae_1 | Clostridium_sensu_stricto_16 |
| Bacteria | Firmicutes | Clostridia | Clostridiales | Clostridiaceae_1 | Clostridium_sensu_stricto_16 |
| Bacteria | Firmicutes | Clostridia | Clostridiales | Clostridiaceae_1 | Clostridium_sensu_stricto_3 |
| Bacteria | Firmicutes | Clostridia | Clostridiales | Clostridiaceae_1 | Clostridium_sensu_stricto_16 |
| Bacteria | Firmicutes | Clostridia | Clostridiales | Clostridiaceae_1 | Clostridium_sensu_stricto_8 |
| Bacteria | Firmicutes | Clostridia | Clostridiales | Ruminococcaceae | NA |
| Bacteria | Firmicutes | Clostridia | Clostridiales | Lachnospiraceae | Anaerocolumna |
| Bacteria | Firmicutes | Clostridia | Clostridiales | Ruminococcaceae | NA |
| Bacteria | Firmicutes | Clostridia | Clostridiales | Family_XI | Finegoldia |
| Bacteria | Firmicutes | Clostridia | Clostridiales | Ruminococcaceae | Hydrogenoanaerobacterium |
| Bacteria | Firmicutes | Clostridia | Clostridiales | Family_XI | Ezakiella |
| Bacteria | Firmicutes | Clostridia | Clostridiales | Lachnospiraceae | Blautia |
| Bacteria | Firmicutes | Clostridia | Clostridiales | Family_XI | Anaerococcus |
| Bacteria | Firmicutes | Clostridia | Clostridiales | Family_XI | Tissierella |
| Bacteria | Firmicutes | Clostridia | Clostridiales | Ruminococcaceae | Faecalibacterium |
| Bacteria | Firmicutes | Clostridia | Clostridiales | Clostridiaceae_2 | Alkaliphilus |
| Bacteria | Firmicutes | Clostridia | Clostridiales | Ruminococcaceae | NA |
| Bacteria | Firmicutes | Clostridia | Clostridiales | Lachnospiraceae | Tyzzerella |
| Bacteria | Firmicutes | Clostridia | Clostridiales | Clostridiaceae_1 | Clostridium_sensu_stricto_1 |
| Bacteria | Proteobacteria | Gammaproteobacteria | Pseudomonadales | Moraxellaceae | Acinetobacter |
| Bacteria | Proteobacteria | Gammaproteobacteria | Betaproteobacteriales | Burkholderiaceae | Alcaligenes |
| Bacteria | Proteobacteria | Gammaproteobacteria | Enterobacteriales | Enterobacteriaceae | NA |
| Bacteria | Proteobacteria | Gammaproteobacteria | Betaproteobacteriales | Burkholderiaceae | Curvibacter |
| Bacteria | Proteobacteria | Gammaproteobacteria | Xanthomonadales | Xanthomonadaceae | Stenotrophomonas |
| Bacteria | Proteobacteria | Gammaproteobacteria | Betaproteobacteriales | Burkholderiaceae | Comamonas |
| Bacteria | Proteobacteria | Gammaproteobacteria | Enterobacteriales | Enterobacteriaceae | Pantoea |
| Bacteria | Proteobacteria | Gammaproteobacteria | Enterobacteriales | Enterobacteriaceae | Proteus |
| Bacteria | Proteobacteria | Gammaproteobacteria | Enterobacteriales | Enterobacteriaceae | Proteus |
| Bacteria | Proteobacteria | Gammaproteobacteria | Enterobacteriales | Enterobacteriaceae | NA |
| Bacteria | Proteobacteria | Gammaproteobacteria | Betaproteobacteriales | Rhodocyclaceae | Azospira |
| Bacteria | Proteobacteria | Gammaproteobacteria | Enterobacteriales | Enterobacteriaceae | Proteus |
| Bacteria | Proteobacteria | Gammaproteobacteria | Betaproteobacteriales | Burkholderiaceae | Comamonas |
| Bacteria | Proteobacteria | Gammaproteobacteria | Enterobacteriales | Enterobacteriaceae | Proteus |
| Bacteria | Proteobacteria | Gammaproteobacteria | Enterobacteriales | Enterobacteriaceae | Proteus |
| Bacteria | Proteobacteria | Gammaproteobacteria | Enterobacteriales | Enterobacteriaceae | Proteus |
| Bacteria | Proteobacteria | Gammaproteobacteria | Enterobacteriales | Enterobacteriaceae | Pectobacterium |
| Bacteria | Proteobacteria | Gammaproteobacteria | Enterobacteriales | Enterobacteriaceae | Proteus |
| Bacteria | Proteobacteria | Gammaproteobacteria | Enterobacteriales | Enterobacteriaceae | Cosenzaea |
| Bacteria | Proteobacteria | Gammaproteobacteria | Enterobacteriales | Enterobacteriaceae | Proteus |
| Bacteria | Proteobacteria | Gammaproteobacteria | Enterobacteriales | Enterobacteriaceae | Cosenzaea |
| Bacteria | Proteobacteria | Gammaproteobacteria | Enterobacteriales | Enterobacteriaceae | NA |
| Bacteria | Proteobacteria | Gammaproteobacteria | Betaproteobacteriales | Burkholderiaceae | Comamonas |
| Bacteria | Proteobacteria | Gammaproteobacteria | Xanthomonadales | Xanthomonadaceae | Stenotrophomonas |
| Bacteria | Proteobacteria | Gammaproteobacteria | Enterobacteriales | Enterobacteriaceae | Pluralibacter |
| Bacteria | Proteobacteria | Gammaproteobacteria | Betaproteobacteriales | Burkholderiaceae | NA |
| Bacteria | Proteobacteria | Gammaproteobacteria | Xanthomonadales | Xanthomonadaceae | Stenotrophomonas |
| Bacteria | Proteobacteria | Gammaproteobacteria | Pseudomonadales | Pseudomonadaceae | Pseudomonas |
| Bacteria | Proteobacteria | Gammaproteobacteria | Xanthomonadales | Xanthomonadaceae | Stenotrophomonas |
| Bacteria | Proteobacteria | Gammaproteobacteria | Xanthomonadales | Xanthomonadaceae | Stenotrophomonas |
| Bacteria | Proteobacteria | Gammaproteobacteria | Betaproteobacteriales | Rhodocyclaceae | NA |
| Bacteria | Proteobacteria | Gammaproteobacteria | Enterobacteriales | Enterobacteriaceae | Morganella |
| Bacteria | Proteobacteria | Gammaproteobacteria | Betaproteobacteriales | Burkholderiaceae | NA |
| Bacteria | Proteobacteria | Gammaproteobacteria | Betaproteobacteriales | Burkholderiaceae | Undibacterium |
| Bacteria | Proteobacteria | Gammaproteobacteria | Enterobacteriales | Enterobacteriaceae | Cosenzaea |
| Bacteria | Proteobacteria | Gammaproteobacteria | Enterobacteriales | Enterobacteriaceae | NA |
| Bacteria | Proteobacteria | Gammaproteobacteria | Enterobacteriales | Enterobacteriaceae | Proteus |
| Bacteria | Proteobacteria | Gammaproteobacteria | Betaproteobacteriales | Chromobacteriaceae | Aquitalea |
| Bacteria | Proteobacteria | Gammaproteobacteria | Betaproteobacteriales | Burkholderiaceae | Delftia |
| Bacteria | Proteobacteria | Gammaproteobacteria | Betaproteobacteriales | Burkholderiaceae | Alcaligenes |
| Bacteria | Proteobacteria | Gammaproteobacteria | Enterobacteriales | Enterobacteriaceae | Cosenzaea |
| Bacteria | Proteobacteria | Gammaproteobacteria | Enterobacteriales | Enterobacteriaceae | Serratia |
| Bacteria | Proteobacteria | Gammaproteobacteria | Betaproteobacteriales | Neisseriaceae | NA |
| Bacteria | Proteobacteria | Gammaproteobacteria | Enterobacteriales | Enterobacteriaceae | Enterobacter |
| Bacteria | Proteobacteria | Gammaproteobacteria | Xanthomonadales | Xanthomonadaceae | Stenotrophomonas |
| Bacteria | Proteobacteria | Gammaproteobacteria | Enterobacteriales | Enterobacteriaceae | Pluralibacter |
| Bacteria | Proteobacteria | Gammaproteobacteria | Betaproteobacteriales | Burkholderiaceae | Achromobacter |
| Bacteria | Proteobacteria | Gammaproteobacteria | Betaproteobacteriales | Burkholderiaceae | Cupriavidus |
| Bacteria | Proteobacteria | Gammaproteobacteria | Betaproteobacteriales | Burkholderiaceae | Comamonas |
| Bacteria | Proteobacteria | Gammaproteobacteria | Enterobacteriales | Enterobacteriaceae | Providencia |
| Bacteria | Proteobacteria | Gammaproteobacteria | Betaproteobacteriales | Burkholderiaceae | Polaromonas |
| Bacteria | Proteobacteria | Gammaproteobacteria | Pseudomonadales | Moraxellaceae | Acinetobacter |
| Bacteria | Proteobacteria | Gammaproteobacteria | Betaproteobacteriales | Burkholderiaceae | Burkholderia-Caballeronia-Paraburkholderia |
| Bacteria | Proteobacteria | Gammaproteobacteria | Betaproteobacteriales | Burkholderiaceae | Curvibacter |
| Bacteria | Proteobacteria | Gammaproteobacteria | Betaproteobacteriales | Burkholderiaceae | Comamonas |
| Bacteria | Proteobacteria | Gammaproteobacteria | Enterobacteriales | Enterobacteriaceae | Morganella |
| Bacteria | Proteobacteria | Gammaproteobacteria | Enterobacteriales | Enterobacteriaceae | Pluralibacter |
| Bacteria | Proteobacteria | Gammaproteobacteria | Salinisphaerales | Solimonadaceae | NA |
| Bacteria | Proteobacteria | Gammaproteobacteria | Gammaproteobacteria_Incertae_Sedis | Unknown_Family | Candidatus_Ovatusbacter |
| Bacteria | Proteobacteria | Gammaproteobacteria | Betaproteobacteriales | Burkholderiaceae | Ralstonia |
| Bacteria | Proteobacteria | Gammaproteobacteria | Pasteurellales | Pasteurellaceae | Haemophilus |
| Bacteria | Proteobacteria | Gammaproteobacteria | Betaproteobacteriales | Burkholderiaceae | Comamonas |
| Bacteria | Proteobacteria | Gammaproteobacteria | Pseudomonadales | Pseudomonadaceae | Pseudomonas |
| Bacteria | Proteobacteria | Gammaproteobacteria | Betaproteobacteriales | Burkholderiaceae | Comamonas |
| Bacteria | Proteobacteria | Gammaproteobacteria | Enterobacteriales | Enterobacteriaceae | Pectobacterium |
| Bacteria | Proteobacteria | Gammaproteobacteria | Betaproteobacteriales | Burkholderiaceae | NA |
| Bacteria | Proteobacteria | Gammaproteobacteria | Enterobacteriales | Enterobacteriaceae | NA |
| Bacteria | Proteobacteria | Gammaproteobacteria | Pseudomonadales | Pseudomonadaceae | Pseudomonas |
| Bacteria | Proteobacteria | Gammaproteobacteria | Betaproteobacteriales | Burkholderiaceae | Aquabacterium |
| Bacteria | Proteobacteria | Gammaproteobacteria | Pseudomonadales | Moraxellaceae | Acinetobacter |
| Bacteria | Proteobacteria | Gammaproteobacteria | Betaproteobacteriales | Burkholderiaceae | Curvibacter |
| Bacteria | Proteobacteria | Gammaproteobacteria | Betaproteobacteriales | Chromobacteriaceae | Aquitalea |
| Bacteria | Proteobacteria | Gammaproteobacteria | Betaproteobacteriales | Burkholderiaceae | Burkholderia-Caballeronia-Paraburkholderia |
| Bacteria | Proteobacteria | Gammaproteobacteria | Betaproteobacteriales | Burkholderiaceae | NA |
| Bacteria | Proteobacteria | Gammaproteobacteria | Enterobacteriales | Enterobacteriaceae | Providencia |
| Bacteria | Proteobacteria | Gammaproteobacteria | Pseudomonadales | Moraxellaceae | Acinetobacter |
| Bacteria | Proteobacteria | Gammaproteobacteria | Pseudomonadales | Moraxellaceae | Acinetobacter |
| Bacteria | Proteobacteria | Gammaproteobacteria | Betaproteobacteriales | Neisseriaceae | Kingella |
| Bacteria | Proteobacteria | Gammaproteobacteria | Xanthomonadales | Rhodanobacteraceae | Luteibacter |
| Bacteria | Proteobacteria | Gammaproteobacteria | Xanthomonadales | Xanthomonadaceae | Pseudoxanthomonas |
| Bacteria | Proteobacteria | Gammaproteobacteria | Betaproteobacteriales | Burkholderiaceae | Massilia |
| Bacteria | Proteobacteria | Gammaproteobacteria | Enterobacteriales | Enterobacteriaceae | Pluralibacter |
| Bacteria | Proteobacteria | Gammaproteobacteria | Xanthomonadales | Xanthomonadaceae | Stenotrophomonas |
| Bacteria | Proteobacteria | Gammaproteobacteria | Alteromonadales | Alteromonadaceae | Alishewanella |
| Bacteria | Proteobacteria | Gammaproteobacteria | Enterobacteriales | Enterobacteriaceae | NA |
| Bacteria | Proteobacteria | Gammaproteobacteria | Enterobacteriales | Enterobacteriaceae | Morganella |
| Bacteria | Proteobacteria | Gammaproteobacteria | Enterobacteriales | Enterobacteriaceae | endosymbionts8 |
| Bacteria | Proteobacteria | Gammaproteobacteria | Betaproteobacteriales | Burkholderiaceae | Comamonas |
| Bacteria | Proteobacteria | Gammaproteobacteria | Salinisphaerales | Solimonadaceae | Nevskia |
| Bacteria | Proteobacteria | Gammaproteobacteria | Betaproteobacteriales | Burkholderiaceae | NA |
| Bacteria | Proteobacteria | Gammaproteobacteria | Betaproteobacteriales | Rhodocyclaceae | Azoarcus |
| Bacteria | Proteobacteria | Gammaproteobacteria | Betaproteobacteriales | A21b | NA |
| Bacteria | Proteobacteria | Gammaproteobacteria | Betaproteobacteriales | Burkholderiaceae | Aquabacterium |
| Bacteria | Proteobacteria | Gammaproteobacteria | Betaproteobacteriales | Burkholderiaceae | Bordetella |
| Bacteria | Proteobacteria | Gammaproteobacteria | Xanthomonadales | Xanthomonadaceae | Stenotrophomonas |
| Bacteria | Proteobacteria | Gammaproteobacteria | Xanthomonadales | Xanthomonadaceae | Thermomonas |
| Bacteria | Proteobacteria | Gammaproteobacteria | Betaproteobacteriales | Burkholderiaceae | Caldimonas |
| Bacteria | Proteobacteria | Gammaproteobacteria | Xanthomonadales | Rhodanobacteraceae | Dokdonella |
| Bacteria | Proteobacteria | Gammaproteobacteria | Betaproteobacteriales | TRA3-20 | NA |
| Bacteria | Proteobacteria | Gammaproteobacteria | Steroidobacterales | Steroidobacteraceae | NA |
| Bacteria | Proteobacteria | Gammaproteobacteria | Xanthomonadales | Xanthomonadaceae | Stenotrophomonas |
| Bacteria | Proteobacteria | Gammaproteobacteria | Xanthomonadales | Xanthomonadaceae | Stenotrophomonas |
| Bacteria | Proteobacteria | Gammaproteobacteria | Betaproteobacteriales | Rhodocyclaceae | Azospira |
| Bacteria | Proteobacteria | Gammaproteobacteria | Pseudomonadales | Moraxellaceae | Acinetobacter |
| Bacteria | Proteobacteria | Gammaproteobacteria | Betaproteobacteriales | Burkholderiaceae | Cupriavidus |
| Bacteria | Proteobacteria | Gammaproteobacteria | Betaproteobacteriales | Burkholderiaceae | Massilia |
| Bacteria | Proteobacteria | Gammaproteobacteria | Betaproteobacteriales | Methylophilaceae | Methylophilus |
| Bacteria | Proteobacteria | Gammaproteobacteria | Betaproteobacteriales | Burkholderiaceae | Cupriavidus |
| Bacteria | Proteobacteria | Gammaproteobacteria | Xanthomonadales | Xanthomonadaceae | Stenotrophomonas |
| Bacteria | Proteobacteria | Gammaproteobacteria | Pseudomonadales | Moraxellaceae | Moraxella |
| Bacteria | Proteobacteria | Gammaproteobacteria | Steroidobacterales | Steroidobacteraceae | NA |
| Bacteria | Proteobacteria | Gammaproteobacteria | Betaproteobacteriales | Burkholderiaceae | Alcaligenes |
| Bacteria | Proteobacteria | Gammaproteobacteria | Pseudomonadales | Pseudomonadaceae | Pseudomonas |
| Bacteria | Proteobacteria | Gammaproteobacteria | Betaproteobacteriales | Burkholderiaceae | Achromobacter |
| Bacteria | Proteobacteria | Gammaproteobacteria | Betaproteobacteriales | Nitrosomonadaceae | MND1 |
| Bacteria | Tenericutes | Mollicutes | Entomoplasmatales | Entomoplasmataceae | Mesoplasma |
| Bacteria | Tenericutes | Mollicutes | Entomoplasmatales | Spiroplasmataceae | Spiroplasma |
| Bacteria | Tenericutes | Mollicutes | Entomoplasmatales | Spiroplasmataceae | Spiroplasma |
