## Supplementary File 1 for "Tissue- and population-level microbiome analysis of the wasp spider *Argiope bruennichi* identifies a novel dominant bacterial symbiont"

Sequence Processing with DADA2


### Sequence Processing with DADA2

###### M.M. Sheffer & M. Bengtsson

###### October 12, 2019

###### Sequence processing was performed according to the dada2 tutorial; find the latest version of the tutorial here.

##### Install dada2

```
if (!requireNamespace("BiocManager", quietly = TRUE))
  install.packages("BiocManager")
BiocManager::install()
BiocManager::install("dada2")
```

```
## package 'dada2' successfully unpacked and MD5 sums checked
## 
## The downloaded binary packages are in
##  C:\Users\Uhl_4\AppData\Local\Temp\Rtmp6pUw90\downloaded_packages
```

```
library(dada2)
```

##### Set working directory and path to fastq files

Remember to change this to the directory where you have downloaded the fastq files (available here)

```
setwd("C:/Users/Uhl_4/Dropbox/Argiope_Microbiome/NewData/fastq")
path<-"C:/Users/Uhl_4/Dropbox/Argiope_Microbiome/NewData/fastq"
fns<-list.files(path)
```

##### Read in fastq files and sort forward and reverse reads

```
fastqs <- fns[grepl(".fastq$", fns)]
fastqs <- sort(fastqs) # Sort ensures forward/reverse reads are in same order
fnFs <- fastqs[grepl("_R1", fastqs)] # Just the forward read files
fnRs <- fastqs[grepl("_R2", fastqs)] # Just the reverse read files
# Get sample names from the first part of the forward read filenames
sample.names <- sapply(strsplit(fnFs, "_"), `[`, 1)
# Fully specify the path for the fnFs and fnRs
fnFs <- file.path(path, fnFs)
fnRs <- file.path(path, fnRs)
```

##### Plot read quality to look for filtering length

```
plotQualityProfile(fnFs[1:2])
```

```
plotQualityProfile(fnRs[1:2])
```

##### Filter according to read quality

```
filt_path <- file.path(path, "filtered") # Place filtered files in filtered/ subdirectory
filtFs <- file.path(filt_path, paste0(sample.names, "_F_filt.fastq.gz"))
filtRs <- file.path(filt_path, paste0(sample.names, "_R_filt.fastq.gz"))
out <- filterAndTrim(fnFs, filtFs, fnRs, filtRs, truncLen=c(200,200),
                     maxN=0, maxEE=c(2,2), truncQ=2, rm.phix=TRUE,
                     compress=TRUE, multithread=FALSE) # On Windows set multithread=FALSE
head(out)
```

```
##                       reads.in reads.out
## 19-1-3C-MS1H_R1.fastq     1805      1500
## 20-1-3D-MS1L_R1.fastq      711       555
## 21-1-3E-MS1P_R1.fastq     4703      3878
## 22-1-3F-MS1S_R1.fastq    32589     26081
## 23-1-3G-MS1M_R1.fastq    24793     22085
## 24-1-3H-MS1F_R1.fastq     5228      4363
```

##### Estimate and plot error rates

```
#estimate error rate
errF <- learnErrors(filtFs, nbases = 1e+08, multithread=TRUE)
```

```
## 115833000 total bases in 579165 reads from 20 samples will be used for learning the error rates.
```

```
errR <- learnErrors(filtRs, nbases = 1e+08, multithread=TRUE)
```

```
## 115833000 total bases in 579165 reads from 20 samples will be used for learning the error rates.
```

```
#plot error rates
plotErrors(errF, nominalQ=FALSE)
```

```
plotErrors(errR, nominalQ=FALSE)
```

##### Dereplicate and infer samples

Dereplication combines identical sequences into unique sequences with corresponding abundance; sample inference applies an algorithm to infer (per sample) the number of reads per unique sequence

```
#dereplication
derepFs <- derepFastq(filtFs, verbose=TRUE)
derepRs <- derepFastq(filtRs, verbose=TRUE)
# Name the derep-class objects by the sample names
names(derepFs) <- sample.names
names(derepRs) <- sample.names

#sample inference
dadaFs <- dada(derepFs, err=errF, multithread=TRUE)
```

```
## Sample 1 - 1500 reads in 619 unique sequences.
## Sample 2 - 555 reads in 253 unique sequences.
## Sample 3 - 3878 reads in 1156 unique sequences.
## Sample 4 - 26081 reads in 8964 unique sequences.
## Sample 5 - 22085 reads in 3605 unique sequences.
## Sample 6 - 4363 reads in 1475 unique sequences.
## Sample 7 - 916 reads in 389 unique sequences.
## Sample 8 - 1814 reads in 783 unique sequences.
## Sample 9 - 1008 reads in 457 unique sequences.
## Sample 10 - 2222 reads in 777 unique sequences.
## Sample 11 - 48043 reads in 7481 unique sequences.
## Sample 12 - 3940 reads in 1475 unique sequences.
## Sample 13 - 27585 reads in 5443 unique sequences.
## Sample 14 - 8868 reads in 3066 unique sequences.
## Sample 15 - 1268 reads in 539 unique sequences.
## Sample 16 - 5748 reads in 2459 unique sequences.
## Sample 17 - 1822 reads in 753 unique sequences.
## Sample 18 - 1298 reads in 477 unique sequences.
## Sample 19 - 158803 reads in 17811 unique sequences.
## Sample 20 - 257368 reads in 22471 unique sequences.
## Sample 21 - 164145 reads in 16413 unique sequences.
## Sample 22 - 165255 reads in 16234 unique sequences.
## Sample 23 - 1021 reads in 439 unique sequences.
## Sample 24 - 3623 reads in 892 unique sequences.
## Sample 25 - 12568 reads in 2121 unique sequences.
## Sample 26 - 27220 reads in 11866 unique sequences.
## Sample 27 - 9510 reads in 2991 unique sequences.
## Sample 28 - 307450 reads in 30922 unique sequences.
## Sample 29 - 847468 reads in 58410 unique sequences.
## Sample 30 - 69534 reads in 11021 unique sequences.
## Sample 31 - 200603 reads in 22383 unique sequences.
## Sample 32 - 67776 reads in 9485 unique sequences.
## Sample 33 - 100134 reads in 13324 unique sequences.
## Sample 34 - 18142 reads in 4279 unique sequences.
## Sample 35 - 12413 reads in 3890 unique sequences.
## Sample 36 - 2145 reads in 875 unique sequences.
## Sample 37 - 8517 reads in 2270 unique sequences.
## Sample 38 - 321961 reads in 29064 unique sequences.
## Sample 39 - 2261 reads in 853 unique sequences.
## Sample 40 - 83521 reads in 10137 unique sequences.
## Sample 41 - 2288 reads in 828 unique sequences.
## Sample 42 - 12442 reads in 2702 unique sequences.
## Sample 43 - 1752 reads in 698 unique sequences.
## Sample 44 - 12768 reads in 3965 unique sequences.
## Sample 45 - 5424 reads in 1847 unique sequences.
## Sample 46 - 25709 reads in 5070 unique sequences.
## Sample 47 - 303065 reads in 29027 unique sequences.
## Sample 48 - 63007 reads in 9204 unique sequences.
## Sample 49 - 435457 reads in 37945 unique sequences.
## Sample 50 - 115360 reads in 12932 unique sequences.
## Sample 51 - 111532 reads in 12769 unique sequences.
## Sample 52 - 16047 reads in 3610 unique sequences.
## Sample 53 - 1956 reads in 812 unique sequences.
## Sample 54 - 6789 reads in 2273 unique sequences.
## Sample 55 - 58970 reads in 6456 unique sequences.
## Sample 56 - 2877 reads in 1099 unique sequences.
## Sample 57 - 128355 reads in 13838 unique sequences.
## Sample 58 - 272812 reads in 23432 unique sequences.
```

```
dadaRs <- dada(derepRs, err=errR, multithread=TRUE)
```

```
## Sample 1 - 1500 reads in 572 unique sequences.
## Sample 2 - 555 reads in 256 unique sequences.
## Sample 3 - 3878 reads in 1125 unique sequences.
## Sample 4 - 26081 reads in 8492 unique sequences.
## Sample 5 - 22085 reads in 3373 unique sequences.
## Sample 6 - 4363 reads in 1419 unique sequences.
## Sample 7 - 916 reads in 388 unique sequences.
## Sample 8 - 1814 reads in 751 unique sequences.
## Sample 9 - 1008 reads in 418 unique sequences.
## Sample 10 - 2222 reads in 702 unique sequences.
## Sample 11 - 48043 reads in 7100 unique sequences.
## Sample 12 - 3940 reads in 1384 unique sequences.
## Sample 13 - 27585 reads in 5024 unique sequences.
## Sample 14 - 8868 reads in 2929 unique sequences.
## Sample 15 - 1268 reads in 504 unique sequences.
## Sample 16 - 5748 reads in 2321 unique sequences.
## Sample 17 - 1822 reads in 749 unique sequences.
## Sample 18 - 1298 reads in 461 unique sequences.
## Sample 19 - 158803 reads in 17536 unique sequences.
## Sample 20 - 257368 reads in 21251 unique sequences.
## Sample 21 - 164145 reads in 15528 unique sequences.
## Sample 22 - 165255 reads in 15267 unique sequences.
## Sample 23 - 1021 reads in 438 unique sequences.
## Sample 24 - 3623 reads in 875 unique sequences.
## Sample 25 - 12568 reads in 2070 unique sequences.
## Sample 26 - 27220 reads in 11017 unique sequences.
## Sample 27 - 9510 reads in 2903 unique sequences.
## Sample 28 - 307450 reads in 29280 unique sequences.
## Sample 29 - 847468 reads in 54553 unique sequences.
## Sample 30 - 69534 reads in 10414 unique sequences.
## Sample 31 - 200603 reads in 21382 unique sequences.
## Sample 32 - 67776 reads in 8860 unique sequences.
## Sample 33 - 100134 reads in 12403 unique sequences.
## Sample 34 - 18142 reads in 4037 unique sequences.
## Sample 35 - 12413 reads in 3741 unique sequences.
## Sample 36 - 2145 reads in 839 unique sequences.
## Sample 37 - 8517 reads in 2184 unique sequences.
## Sample 38 - 321961 reads in 27002 unique sequences.
## Sample 39 - 2261 reads in 799 unique sequences.
## Sample 40 - 83521 reads in 9572 unique sequences.
## Sample 41 - 2288 reads in 791 unique sequences.
## Sample 42 - 12442 reads in 2464 unique sequences.
## Sample 43 - 1752 reads in 698 unique sequences.
## Sample 44 - 12768 reads in 3677 unique sequences.
## Sample 45 - 5424 reads in 1703 unique sequences.
## Sample 46 - 25709 reads in 4672 unique sequences.
## Sample 47 - 303065 reads in 28099 unique sequences.
## Sample 48 - 63007 reads in 8610 unique sequences.
## Sample 49 - 435457 reads in 35744 unique sequences.
## Sample 50 - 115360 reads in 11549 unique sequences.
## Sample 51 - 111532 reads in 12027 unique sequences.
## Sample 52 - 16047 reads in 3446 unique sequences.
## Sample 53 - 1956 reads in 737 unique sequences.
## Sample 54 - 6789 reads in 2120 unique sequences.
## Sample 55 - 58970 reads in 6174 unique sequences.
## Sample 56 - 2877 reads in 1073 unique sequences.
## Sample 57 - 128355 reads in 12974 unique sequences.
## Sample 58 - 272812 reads in 22116 unique sequences.
```

##### Merge forward and reverse reads, construct sequence table

```
mergers <- mergePairs(dadaFs, derepFs, dadaRs, derepRs, verbose=TRUE)
# Inspect the merger data.frame from the first sample
head(mergers[[1]])
```

```
##                                                                                                                                                                                                                                                        sequence
## 1                           TACAAGGAAGACTAGTGTTATTCATCTTAATTAGGTTTAAAGGGTACCTAGACAGTATTTCTAGCCTCAAAAGGGAACAGACTTACTAGAGTTTTATGGGAGAGGAAAATATTAGAACCATTGGAGTAGAGATAAAATGTTTTGATACTAATGGGACGGATAGCGGCGAAGGCAAACCTCTATGTAATAACTGACGTTGAGGGACGAAGGCTTGGGGAGCGAATAGG
## 2 TACGTAGGGTGCAAGCGTTAATCGGAATTACTGGGCGTAAAGCGTGCGCAGGCGGTTATGCAAGACAGAGGTGAAATCCCCGGGCTCAACCTGGGAACTGCCTTTGTGACTGCATAGCTAGAGTACGGTAGAGGGGGATGGAATTCCGCGTGTAGCAGTGAAATGCGTAGATATGCGGAGGAACACCGATGGCGAAGGCAATCCCCTGGACCTGTACTGACGCTCATGCACGAAAGCGTGGGGAGCAAACAGG
## 3 TACAGAGGGTGCAAGCGTTAATCGGAATTACTGGGCGTAAAGCGTGCGTAGACGGTTACATAAGTCGGGTGTGAAAGCCCCGGGCTCAACCTGGGAATTGCATTCGAGACTGCGTAGCTAGGGTGCGGAAGAGGGAAGCGGAATTTCCGGTGTAGCGGTGAAATGCGTAGATATCGGAAGGAACACCAGTGGCGAAAGCGGCTTCCTGGTCCAGCACCGACGTTCAGGCACGAAAGCGTGGGGAGCAAACAGG
## 4 TACGAAGGGGGCTAGCGTTGCTCGGAATTACTGGGCGTAAAGGGCGCGTAGGCGGACAGTTAAGTTGGGGGTGAAAGCCCGGGGCTCAACCTCGGAATTGCCTTCAATACTGGCTGTCTTGAGTACGGGAGAGGTGAGTGGAACTCCGAGTGTAGAGGTGAAATTCGTAGATATTCGGAAGAACACCAGTGGCGAAGGCGACTCACTGGCCCGTTACTGACGCTGAGGCGCGAAAGCGTGGGGAGCAAACAGG
## 5  TACGTAGGTGGCAAGCGTTATCCGGAATTATTGGGCGTAAAGAGTGAGCAGGCGGTTTTTTAAGTTTAATGTTAAAGGTTAAGACTCAATTTTAATTCGCATTAAAAACTGGAAAACTAGAGTGTGGTAGAGGATAATGGAATTCTGTATGTAGTGGTGAAATACGTAGATATACGGAGGAACATCAATTGCGAAGGCAATTATCTGGACCATTACTGACGCTCAGTCACGAAAGCGTGGGGAGCAAACTGG
## 6 TACGGAGGGTGCAAGCGTTAATCGGAATTACTGGGCGTAAAGCGCACGCAGGTGGCTGAGTCAGTCGATTGTGAAAGCCCTGGGCTTAACCTGGGAATTGCAGTCGATACTACTCAGCTAGAGTATGGGAGAGGGCAGTGGAATTCCCGGTGTAGCGGTGAAATGCGTAGATATCGGGAGGAACATCAGTGGCGAAGGCGGCTGCCTGGCCCAATACTGACACTCAGGTGCGACAGCGTGGGGAGCAAACAGG
##   abundance forward reverse nmatch nmismatch nindel prefer accept
## 1       534       1       1    173         0      0      2   TRUE
## 2       149       2       2    147         0      0      2   TRUE
## 3        95       3       3    147         0      0      2   TRUE
## 4        71       5       4    147         0      0      2   TRUE
## 5        66       4       5    148         0      0      1   TRUE
## 6        53       6       6    147         0      0      2   TRUE
```

```
#construct sequence table
seqtab<-makeSequenceTable(mergers)
dim(seqtab)
```

```
## [1]   58 2246
```

```
#inspect distribution of sequence lengths
table(nchar(getSequences(seqtab)))
```

```
## 
##  203  215  220  221  222  223  224  226  227  228  233  234  236  237  238 
##    1    1    5    1    1   11    2    2   12    2    1    1    1    1    1 
##  242  244  245  246  249  250  251  252  253  254  255  256  257  264  265 
##    1    1    2    1    1    1    6   84 1938  151    5    1    2    1    1 
##  272  274  278  285  290 
##    2    1    2    1    1
```

```
#remove sequences that are too long/too short
seqtab2 <- seqtab[,nchar(colnames(seqtab)) %in% seq(251,254)]
```

##### Remove chimeras

```
seqtab.nochim <- removeBimeraDenovo(seqtab2, method="consensus", multithread=TRUE, verbose=TRUE)
```

```
## Identified 29 bimeras out of 2179 input sequences.
```

```
dim(seqtab.nochim)
```

```
## [1]   58 2150
```

```
sum(seqtab.nochim)/sum(seqtab) #percent of non-chimeras in sequence reads
```

```
## [1] 0.99772
```

##### Taxonomic assignment (use latest Silva download from tutorial)

```
taxa <- assignTaxonomy(seqtab.nochim, "C:/Users/Uhl_4/Dropbox/Argiope_Microbiome/NewData/fastq/silva_nr_v132_train_set.fa.gz", multithread=TRUE) #change file path to latest Silva download
taxa.print <- taxa # Removing sequence rownames for display only
rownames(taxa.print) <- NULL
head(taxa.print)
```

```
##      Kingdom    Phylum           Class                
## [1,] "Bacteria" NA               NA                   
## [2,] "Bacteria" "Tenericutes"    "Mollicutes"         
## [3,] "Bacteria" "Proteobacteria" "Alphaproteobacteria"
## [4,] "Bacteria" "Proteobacteria" "Alphaproteobacteria"
## [5,] "Bacteria" "Proteobacteria" "Gammaproteobacteria"
## [6,] "Bacteria" "Proteobacteria" "Gammaproteobacteria"
##      Order                                Family              
## [1,] NA                                   NA                  
## [2,] "Entomoplasmatales"                  "Entomoplasmataceae"
## [3,] "Caulobacterales"                    "Caulobacteraceae"  
## [4,] "Rhizobiales"                        "Beijerinckiaceae"  
## [5,] "Gammaproteobacteria_Incertae_Sedis" "Unknown_Family"    
## [6,] "Pseudomonadales"                    "Moraxellaceae"     
##      Genus          
## [1,] NA             
## [2,] "Mesoplasma"   
## [3,] NA             
## [4,] NA             
## [5,] "Acidibacter"  
## [6,] "Acinetobacter"
```

##### Write sequence abundance table and taxonomic assignment table

```
library(ape)

seq<-as.data.frame(seqtab.nochim)
fasta<-write.dna(matrix(colnames(seq)), file=file.path(path, "argiope_seqs.fasta"), format="fasta", colw=max(nchar(colnames(seq)))) 

write.csv(seq,file=file.path(path, "argiope_ASVseqtab.csv") ) 
write.csv(taxa,file=file.path(path, "argiope_ASVtaxonomy.csv"))
```
