## Supplementary File 2 for "Tissue- and population-level microbiome analysis of the wasp spider *Argiope bruennichi* identifies a novel dominant bacterial symbiont"

Argiope Microbiome Data Analysis


### Argiope Microbiome Data Analysis

###### M.M. Sheffer & M. Bengtsson

###### October 12, 2019

##### Install and load packages

```
install.packages("vegan",repos="https://cloud.r-project.org")
```

```
## package 'vegan' successfully unpacked and MD5 sums checked
## 
## The downloaded binary packages are in
##  C:\Users\Uhl_4\AppData\Local\Temp\Rtmp0yyiVp\downloaded_packages
```

```
library(vegan)

install.packages("ggplot2",repos="https://cloud.r-project.org")
```

```
## package 'ggplot2' successfully unpacked and MD5 sums checked
## 
## The downloaded binary packages are in
##  C:\Users\Uhl_4\AppData\Local\Temp\Rtmp0yyiVp\downloaded_packages
```

```
library(ggplot2)
```

##### Read in data files

Remember to change this to the directory where you have downloaded the data files (available here)

```
#read in OTU table
seq<-read.csv("C:/Users/Uhl_4/Dropbox/Argiope_Microbiome/R-data/argiope_seq.csv", header=T, row.names = 1)
colnames(seq)<-1:ncol(seq)

#read in sample meta data
meta<-read.csv("C:/Users/Uhl_4/Dropbox/Argiope_Microbiome/R-data/argiope_meta.csv", header=T, row.names = 1)

#read in taxonomic assignment
tax<-read.csv("C:/Users/Uhl_4/Dropbox/Argiope_Microbiome/R-data/argiope_prelim_tax.csv", header=T, row.names = 1)
rownames(tax)<-1:ncol(seq)
```

##### Filter data

We filtered the data through several cutoffs, based on sequencing depth and presence in negative controls

```
#calculate sequencing depth
meta$libsize<-rowSums(seq)

#cut out samples with low sequencing depth
meta1<-meta[meta$libsize>4000,] 
seq1<-seq[meta$libsize>4000,] 

#cut out sequences with zero abundance
seltax<-which(colSums(seq1)>0)
seq2<-seq1[,seltax]
tax1<-tax[seltax,]

#calculate abundance of sequences in negative control samples
tax1$Sum<-colSums(seq2)
tax1$Sum_control<-colSums(seq2[meta1$Tissue=="control",])

#cut out sequences with high abundance in control samples
seltax<-which(tax1$Sum_control<50)
seq3<-seq2[,seltax]
tax2<-tax1[seltax,]

#cut out control samples
meta2<-meta1[meta1$Tissue!="control",] 
seq4<-seq3[meta1$Tissue!="control",] 

#cut out sequences with zero abundance
seltax<-which(colSums(seq4)>0)
seq5<-seq4[,seltax]
tax3<-tax2[seltax,]

#observed sequence variants
meta2$nseq<-apply(seq5>0, 1, sum)
meta2$libsize1<-rowSums(seq5)

#cut out samples with low sequencing depth again
meta3<-meta2[meta2$libsize1>400,] 
seq6<-seq5[meta2$libsize1>400,] 

#cut out sequences with zero abundance
seltax<-which(colSums(seq5)>0)
seq7<-seq6[,seltax]
tax4<-tax3[seltax,]

#observed sequence variants
meta3$nseq1<-apply(seq7>0, 1, sum)
meta3$libsize2<-rowSums(seq7)

min(meta3$libsize2)
```

```
## [1] 477
```

```
max(meta3$libsize2)
```

```
## [1] 629137
```

```
mean(meta3$libsize2)
```

```
## [1] 41182.67
```

```
sum(meta3$libsize2)
```

```
## [1] 1770855
```

#### Pie charts

##### Data aggregation

We have one amplicon sequence variant (ASV) that is highly abundant, but has no taxonomic assignment. Therefore, we name it “Unknown Symbiont” so that it remains separate during aggregation

```
tax4[,2]<-as.character(tax4[,2])
tax4[1,2]<-"Unknown Symbiont"
tax4[,3]<-as.character(tax4[,3])
tax4[1,3]<-"Unknown Symbiont"
tax4[,4]<-as.character(tax4[,4])
tax4[1,4]<-"Unknown Symbiont"
tax4[,5]<-as.character(tax4[,5])
tax4[1,5]<-"Unknown Symbiont"
tax4[,6]<-as.character(tax4[,6])
tax4[1,6]<-"Unknown Symbiont"
```

Now, we replace all other NAs in the taxon table with “Unclassified”" category (different from Unknown Symbiont category, which represents just one ASV)

```
for(i in 2:6) {
  tax4[,i][is.na(tax4[,i])] <- "Unclassified"
}
```

Aggregate data to class level for pie charts

```
agg<-aggregate(t(seq7), list(class=tax4$Class), sum)
ix<-as.character(agg[,1])
class<-data.frame(t(agg[,-1]))
names(class)<-ix

#Make an 'other' category for classes with low abundance
Other<-class[,colSums(class)<1000]
class1<-class[,colSums(class)>1000]
class1$Other<-rowSums(Other)
```

##### Subset data by tissue type and population

```
bl_g<-as.data.frame(t(class1[which(meta3$Country=="germany"&meta3$Tissue=="book_lung"),]))
f_g<-as.data.frame(t(class1[which(meta3$Country=="germany"&meta3$Tissue=="fecal"),]))
h_g<-as.data.frame(t(class1[which(meta3$Country=="germany"&meta3$Tissue=="hemolymph"),]))
l_g<-as.data.frame(t(class1[which(meta3$Country=="germany"&meta3$Tissue=="leg"),]))
m_g<-as.data.frame(t(class1[which(meta3$Country=="germany"&meta3$Tissue=="midgut"),]))
o_g<-as.data.frame(t(class1[which(meta3$Country=="germany"&meta3$Tissue=="ovary"),]))
es_g<-as.data.frame(t(class1[which(meta3$Country=="germany"&meta3$Tissue=="egg_sac"),]))
sp_g<-as.data.frame(t(class1[which(meta3$Country=="germany"&meta3$Tissue=="spiderling"),]))
p_g<-as.data.frame(t(class1[which(meta3$Country=="germany"&meta3$Tissue=="prosoma"),]))
sg_g<-as.data.frame(t(class1[which(meta3$Country=="germany"&meta3$Tissue=="silk_gland"),]))


bl_e<-as.data.frame(t(class1[which(meta3$Country=="estonia"&meta3$Tissue=="book_lung"),]))
f_e<-as.data.frame(t(class1[which(meta3$Country=="estonia"&meta3$Tissue=="fecal"),]))
h_e<-as.data.frame(t(class1[which(meta3$Country=="estonia"&meta3$Tissue=="hemolymph"),]))
l_e<-as.data.frame(t(class1[which(meta3$Country=="estonia"&meta3$Tissue=="leg"),]))
m_e<-as.data.frame(t(class1[which(meta3$Country=="estonia"&meta3$Tissue=="midgut"),]))
o_e<-as.data.frame(t(class1[which(meta3$Country=="estonia"&meta3$Tissue=="ovary"),]))
es_e<-as.data.frame(t(class1[which(meta3$Country=="estonia"&meta3$Tissue=="egg_sac"),]))
sp_e<-as.data.frame(t(class1[which(meta3$Country=="estonia"&meta3$Tissue=="spiderling"),]))
p_e<-as.data.frame(t(class1[which(meta3$Country=="estonia"&meta3$Tissue=="prosoma"),]))
sg_e<-as.data.frame(t(class1[which(meta3$Country=="estonia"&meta3$Tissue=="silk_gland"),]))
```

##### Draw pie charts for each tissue type per country

Note: in practice, we wrote these plots to a PDF, and then arranged them in the figure with color coding for tissue type in a vector graphics editor. Here, the plots are in their original output.

```
#create a vector of 9 distinct colors####
distinct<-c('#e6194b', '#3cb44b', '#ffe119', '#bcf60c', '#f58231', '#911eb4', '#46f0f0', '#e6beff', '#4363d8')

##booklungs
#germany
classnames<-rownames(bl_g)
bl_g_sum<-rowSums(bl_g)
bl_g_final<-data.frame(class = classnames,sum=bl_g_sum)
bpbl_g<-ggplot(bl_g_final,aes(x="Germany Book Lungs",y=sum,fill=class))+
  geom_bar(width=1,stat="identity")
pieBL_g<-bpbl_g+coord_polar("y",start=0)
pieBL_g+scale_fill_manual(values=distinct)+theme(legend.title =element_text(size=7), legend.text = element_text(size=5), legend.key.size = unit(.2,"cm"))
#estonia
classnames<-rownames(bl_e)
bl_e_sum<-rowSums(bl_e)
bl_e_final<-data.frame(class = classnames,sum=bl_e_sum)
bpbl_e<-ggplot(bl_e_final,aes(x="Estonia Book Lungs",y=sum,fill=class))+
  geom_bar(width=1,stat="identity")
pieBL_e<-bpbl_e+coord_polar("y",start=0)
pieBL_e+scale_fill_manual(values=distinct)+theme(legend.title =element_text(size=7), legend.text = element_text(size=5), legend.key.size = unit(.2,"cm"))

##fecal pellets
#germany
classnames<-rownames(f_g)
f_g_sum<-rowSums(f_g)
f_g_final<-data.frame(class = classnames,sum=f_g_sum)
bpf_g<-ggplot(f_g_final,aes(x="Germany Fecal Pellets",y=sum,fill=class))+
  geom_bar(width=1,stat="identity")
pieF_g<-bpf_g+coord_polar("y",start=0)
pieF_g+scale_fill_manual(values=distinct)+theme(legend.title =element_text(size=7), legend.text = element_text(size=5), legend.key.size = unit(.2,"cm"))
#estonia
classnames<-rownames(f_e)
f_e_sum<-rowSums(f_e)
f_e_final<-data.frame(class = classnames,sum=f_e_sum)
bpf_e<-ggplot(f_e_final,aes(x="Estonia Fecal Pellets",y=sum,fill=class))+
  geom_bar(width=1,stat="identity")
pieF_e<-bpf_e+coord_polar("y",start=0)
pieF_e+scale_fill_manual(values=distinct)+theme(legend.title =element_text(size=7), legend.text = element_text(size=5), legend.key.size = unit(.2,"cm"))

##hemolymph
#germany
classnames<-rownames(h_g)
h_g_sum<-rowSums(h_g)
h_g_final<-data.frame(class = classnames,sum=h_g_sum)
bph_g<-ggplot(h_g_final,aes(x="Germany Hemolymph",y=sum,fill=class))+
  geom_bar(width=1,stat="identity")
pieH_g<-bph_g+coord_polar("y",start=0)
pieH_g+scale_fill_manual(values=distinct)+theme(legend.title =element_text(size=7), legend.text = element_text(size=5), legend.key.size = unit(.2,"cm"))
#estonia
classnames<-rownames(h_e)
h_e_sum<-rowSums(h_e)
h_e_final<-data.frame(class = classnames,sum=h_e_sum)
bph_e<-ggplot(h_e_final,aes(x="Estonia Hemolymph",y=sum,fill=class))+
  geom_bar(width=1,stat="identity")
pieH_e<-bph_e+coord_polar("y",start=0)
pieH_e+scale_fill_manual(values=distinct)+theme(legend.title =element_text(size=7), legend.text = element_text(size=5), legend.key.size = unit(.2,"cm"))

##legs
#germany
classnames<-rownames(l_g)
l_g_sum<-rowSums(l_g)
l_g_final<-data.frame(class = classnames,sum=l_g_sum)
bpl_g<-ggplot(l_g_final,aes(x="Germany Leg",y=sum,fill=class))+
  geom_bar(width=1,stat="identity")
pieL_g<-bpl_g+coord_polar("y",start=0)
pieL_g+scale_fill_manual(values=distinct)+theme(legend.title =element_text(size=7), legend.text = element_text(size=5), legend.key.size = unit(.2,"cm"))
#estonia
classnames<-rownames(l_e)
l_e_sum<-rowSums(l_e)
l_e_final<-data.frame(class = classnames,sum=l_e_sum)
bpl_e<-ggplot(l_e_final,aes(x="Estonia Leg",y=sum,fill=class))+
  geom_bar(width=1,stat="identity")
pieL_e<-bpl_e+coord_polar("y",start=0)
pieL_e+scale_fill_manual(values=distinct)+theme(legend.title =element_text(size=7), legend.text = element_text(size=5), legend.key.size = unit(.2,"cm"))

##midgut
#germany
classnames<-rownames(m_g)
m_g_sum<-rowSums(m_g)
m_g_final<-data.frame(class = classnames,sum=m_g_sum)
bpm_g<-ggplot(m_g_final,aes(x="Germany Midgut",y=sum,fill=class))+
  geom_bar(width=1,stat="identity")
pieM_g<-bpm_g+coord_polar("y",start=0)
pieM_g+scale_fill_manual(values=distinct)+theme(legend.title =element_text(size=7), legend.text = element_text(size=5), legend.key.size = unit(.2,"cm"))
#estonia
classnames<-rownames(m_e)
m_e_sum<-rowSums(m_e)
m_e_final<-data.frame(class = classnames,sum=m_e_sum)
bpm_e<-ggplot(m_e_final,aes(x="Estonia Midgut",y=sum,fill=class))+
  geom_bar(width=1,stat="identity")
pieM_e<-bpm_e+coord_polar("y",start=0)
pieM_e+scale_fill_manual(values=distinct)+theme(legend.title =element_text(size=7), legend.text = element_text(size=5), legend.key.size = unit(.2,"cm"))

##ovaries
#germany
classnames<-rownames(o_g)
o_g_sum<-rowSums(o_g)
o_g_final<-data.frame(class = classnames,sum=o_g_sum)
bpo_g<-ggplot(o_g_final,aes(x="Germany Ovary",y=sum,fill=class))+
  geom_bar(width=1,stat="identity")
pieO_g<-bpo_g+coord_polar("y",start=0)
pieO_g+scale_fill_manual(values=distinct)+theme(legend.title =element_text(size=7), legend.text = element_text(size=5), legend.key.size = unit(.2,"cm"))
#estonia
classnames<-rownames(o_e)
o_e_sum<-rowSums(o_e)
o_e_final<-data.frame(class = classnames,sum=o_e_sum)
bpo_e<-ggplot(o_e_final,aes(x="Estonia Ovary",y=sum,fill=class))+
  geom_bar(width=1,stat="identity")
pieO_e<-bpo_e+coord_polar("y",start=0)
pieO_e+scale_fill_manual(values=distinct)+theme(legend.title =element_text(size=7), legend.text = element_text(size=5), legend.key.size = unit(.2,"cm"))

##spiderlings
#germany
classnames<-rownames(sp_g)
sp_g_sum<-rowSums(sp_g)
sp_g_final<-data.frame(class = classnames,sum=sp_g_sum)
bpsp_g<-ggplot(sp_g_final,aes(x="Germany Spiderlings",y=sum,fill=class))+
  geom_bar(width=1,stat="identity")
piesp_g<-bpsp_g+coord_polar("y",start=0)
piesp_g+scale_fill_manual(values=distinct)+theme(legend.title =element_text(size=7), legend.text = element_text(size=5), legend.key.size = unit(.2,"cm"))
#estonia
classnames<-rownames(sp_e)
sp_e_sum<-rowSums(sp_e)
sp_e_final<-data.frame(class = classnames,sum=sp_e_sum)
bpsp_e<-ggplot(sp_e_final,aes(x="Estonia Spiderlings",y=sum,fill=class))+
  geom_bar(width=1,stat="identity")
piesp_e<-bpsp_e+coord_polar("y",start=0)
piesp_e+scale_fill_manual(values=distinct)+theme(legend.title =element_text(size=7), legend.text = element_text(size=5), legend.key.size = unit(.2,"cm"))

##prosoma
#germany
classnames<-rownames(p_g)
p_g_sum<-rowSums(p_g)
p_g_final<-data.frame(class = classnames,sum=p_g_sum)
bpp_g<-ggplot(p_g_final,aes(x="Germany Prosoma",y=sum,fill=class))+
  geom_bar(width=1,stat="identity")
pieP_g<-bpp_g+coord_polar("y",start=0)
pieP_g+scale_fill_manual(values=distinct)+theme(legend.title =element_text(size=7), legend.text = element_text(size=5), legend.key.size = unit(.2,"cm"))
#estonia
classnames<-rownames(p_e)
p_e_sum<-rowSums(p_e)
p_e_final<-data.frame(class = classnames,sum=p_e_sum)
bpp_e<-ggplot(p_e_final,aes(x="Estonia Prosoma",y=sum,fill=class))+
  geom_bar(width=1,stat="identity")
pieP_e<-bpp_e+coord_polar("y",start=0)
pieP_e+scale_fill_manual(values=distinct)+theme(legend.title =element_text(size=7), legend.text = element_text(size=5), legend.key.size = unit(.2,"cm"))

##silk glands
#germany
classnames<-rownames(sg_g)
sg_g_sum<-rowSums(sg_g)
sg_g_final<-data.frame(class = classnames,sum=sg_g_sum)
bpsg_g<-ggplot(sg_g_final,aes(x="Germany Silk Gland",y=sum,fill=class))+
  geom_bar(width=1,stat="identity")
pieSG_g<-bpsg_g+coord_polar("y",start=0)
pieSG_g+scale_fill_manual(values=distinct)+theme(legend.title =element_text(size=7), legend.text = element_text(size=5), legend.key.size = unit(.2,"cm"))
#estonia
classnames<-rownames(sg_e)
sg_e_sum<-rowSums(sg_e)
sg_e_final<-data.frame(class = classnames,sum=sg_e_sum)
bpsg_e<-ggplot(sg_e_final,aes(x="Estonia Silk Gland",y=sum,fill=class))+
  geom_bar(width=1,stat="identity")
pieSG_e<-bpsg_e+coord_polar("y",start=0)
pieSG_e+scale_fill_manual(values=distinct)+theme(legend.title =element_text(size=7), legend.text = element_text(size=5), legend.key.size = unit(.2,"cm"))
```

#### PERMANOVA and nMDS

##### Remove spiderlings and unknown symbiont

Note: in this analysis, individuals 7 and 8 (spiderlings) should be excluded. We also performed the analysis with and without the most dominant symbiont. The next few lines of code remove individuals 7 and 8 in both cases, and in one case remove the dominant symbiont

```
#remove only 7 and 8, not unknown
seq10<-seq7[1:39,]
meta5<-meta3[1:39,]

#remove unknown symbiont and individuals 7 and 8
seq8<-seq7[1:39,2:574]
tax5<-tax4[2:574,]
meta4<-meta3[1:39,]
```

##### Transform data and run PERMANOVA analysis

```
#hellinger transform dataset with everything (spiderlings and unknown symbiont) removed, then run PERMANOVA
seq11<-decostand(seq8, method = "hellinger")
mds1<-metaMDS(seq11) #MDS
```

```
## Run 0 stress 0.1430268 
## Run 1 stress 0.1568225 
## Run 2 stress 0.1448709 
## Run 3 stress 0.1436025 
## Run 4 stress 0.1498635 
## Run 5 stress 0.1512936 
## Run 6 stress 0.1431038 
## ... Procrustes: rmse 0.01331094  max resid 0.05010705 
## Run 7 stress 0.1501077 
## Run 8 stress 0.1569017 
## Run 9 stress 0.1466693 
## Run 10 stress 0.1525403 
## Run 11 stress 0.1517435 
## Run 12 stress 0.1521703 
## Run 13 stress 0.1592237 
## Run 14 stress 0.1567277 
## Run 15 stress 0.1518873 
## Run 16 stress 0.1458019 
## Run 17 stress 0.1539412 
## Run 18 stress 0.1479869 
## Run 19 stress 0.1464997 
## Run 20 stress 0.154332 
## *** No convergence -- monoMDS stopping criteria:
##      5: no. of iterations >= maxit
##     15: stress ratio > sratmax
```

```
adonis(seq11~Tissue*Country*Individual,data=meta4) #PERMANOVA
```

```
## 
## Call:
## adonis(formula = seq11 ~ Tissue * Country * Individual, data = meta4) 
## 
## Permutation: free
## Number of permutations: 999
## 
## Terms added sequentially (first to last)
## 
##                           Df SumsOfSqs MeanSqs F.Model      R2 Pr(>F)    
## Tissue                     7    3.2281 0.46115 1.01792 0.18034  0.375    
## Country                    1    0.8095 0.80948 1.78680 0.04522  0.004 ** 
## Individual                 1    1.0551 1.05510 2.32896 0.05895  0.001 ***
## Tissue:Country             7    3.1098 0.44426 0.98063 0.17374  0.583    
## Tissue:Individual          7    2.7430 0.39185 0.86495 0.15324  0.949    
## Country:Individual         1    0.7817 0.78173 1.72556 0.04367  0.007 ** 
## Tissue:Country:Individual  5    2.0951 0.41903 0.92493 0.11705  0.810    
## Residuals                  9    4.0773 0.45303         0.22779           
## Total                     38   17.8996                 1.00000           
## ---
## Signif. codes:  0 '***' 0.001 '**' 0.01 '*' 0.05 '.' 0.1 ' ' 1
```

```
#hellinger transform dataset with only spiderlings removed, then run PERMANOVA
seq13<-decostand(seq10, method = "hellinger")
mds3<-metaMDS(seq13) #MDS
```

```
## Run 0 stress 0.09848903 
## Run 1 stress 0.1022007 
## Run 2 stress 0.09553119 
## ... New best solution
## ... Procrustes: rmse 0.03999055  max resid 0.1246627 
## Run 3 stress 0.1024887 
## Run 4 stress 0.0955555 
## ... Procrustes: rmse 0.07423228  max resid 0.2245742 
## Run 5 stress 0.1095877 
## Run 6 stress 0.09582217 
## ... Procrustes: rmse 0.06971944  max resid 0.2190002 
## Run 7 stress 0.1023689 
## Run 8 stress 0.1005559 
## Run 9 stress 0.1002051 
## Run 10 stress 0.09628609 
## Run 11 stress 0.09561395 
## ... Procrustes: rmse 0.04751067  max resid 0.2438851 
## Run 12 stress 0.09966884 
## Run 13 stress 0.1107935 
## Run 14 stress 0.095648 
## ... Procrustes: rmse 0.01775467  max resid 0.08201987 
## Run 15 stress 0.09630224 
## Run 16 stress 0.1107173 
## Run 17 stress 0.09485696 
## ... New best solution
## ... Procrustes: rmse 0.07708131  max resid 0.2488657 
## Run 18 stress 0.09410852 
## ... New best solution
## ... Procrustes: rmse 0.01661928  max resid 0.0501142 
## Run 19 stress 0.0967829 
## Run 20 stress 0.1105897 
## *** No convergence -- monoMDS stopping criteria:
##      7: no. of iterations >= maxit
##     13: stress ratio > sratmax
```

```
adonis(seq13~Tissue*Country*Individual,data=meta5) #PERMANOVA
```

```
## 
## Call:
## adonis(formula = seq13 ~ Tissue * Country * Individual, data = meta5) 
## 
## Permutation: free
## Number of permutations: 999
## 
## Terms added sequentially (first to last)
## 
##                           Df SumsOfSqs MeanSqs F.Model      R2 Pr(>F)  
## Tissue                     7    1.7071 0.24387 1.36265 0.23140  0.118  
## Country                    1    0.2912 0.29116 1.62687 0.03947  0.083 .
## Individual                 1    0.2957 0.29574 1.65247 0.04009  0.084 .
## Tissue:Country             7    1.6046 0.22923 1.28082 0.21751  0.192  
## Tissue:Individual          7    0.8916 0.12737 0.71168 0.12086  0.844  
## Country:Individual         1    0.4210 0.42101 2.35237 0.05707  0.014 *
## Tissue:Country:Individual  5    0.5553 0.11107 0.62060 0.07528  0.915  
## Residuals                  9    1.6107 0.17897         0.21834         
## Total                     38    7.3773                 1.00000         
## ---
## Signif. codes:  0 '***' 0.001 '**' 0.01 '*' 0.05 '.' 0.1 ' ' 1
```

Note: the results do not change much with or without the unknown symbiont; just the level of significance changes. Therefore, we only plot one of the groupings.

##### Visualize data with nMDS plots

Note: because the analysis is permutation based, this graph will change each time it is calculated. Therefore, the generated plot in this document, and any plots of yours if you choose to replicate the analysis on your own, will not be exactly the same as the figure in the publication.

```
##create vectors of distinct colors for each grouping variable, shapes for tissue type
levels(droplevels(meta4$Tissue))
```

```
## [1] "book_lung"  "fecal"      "hemolymph"  "leg"        "midgut"    
## [6] "ovary"      "prosoma"    "silk_gland"
```

```
tissueShapes<-c(21,22,23,24,25,3,4,8)

levels(meta4$Country)
```

```
## [1] "estonia" "germany"
```

```
countrycol<-c('#ffe119', '#4363d8')

meta4$Individual<-as.factor(meta4$Individual)
levels(droplevels(meta4$Individual))
```

```
## [1] "1" "2" "3" "4" "5" "6"
```

```
indcols<-c("#ffe119","#fff7bf","#a69100",'#4363d8',"#bcc9f8","#061b68")

##plot with unknown symbiont excluded
plot(mds1, display="sites", type = "none")
points(mds1, col=indcols[droplevels(meta4$Individual)], bg=indcols[droplevels(meta4$Individual)],pch=tissueShapes[droplevels(meta4$Tissue)], cex=2)
legend("topright",legend=c("Estonia 1","Estonia 2","Estonia 3","Germany 1","Germany 2","Germany 3"),fill=indcols,ncol=2,cex=.8)
ordiellipse(mds1,droplevels(meta4$Individual),display="sites",draw="polygon",col=indcols,alpha=75,border=c("#a69100","#a69100","#a69100","#061b68","#061b68","#061b68"),kind = "se",conf=.99)
legend("topleft",legend=c("Book Lungs","Fecal Pellets","Hemolymph","Leg","Midgut","Ovaries","Prosoma","Silk Glands"),pch =tissueShapes ,ncol=1,cex=.8)
```
