## Supplementary Table 2 for "Tissue- and population-level microbiome analysis of the wasp spider *Argiope bruennichi* identifies a novel dominant bacterial symbiont"

**Additional File 4: Accession Numbers for Gene Tree**

A table detailing all of the sequences downloaded from Silva and GenBank used to generate a 16S rRNA gene tree

| **Accession Number** | **Organism Name** | **Phylum** | **Database** |
| --- | --- | --- | --- |
| AY327470 | *Cardinium* endosymbiont of *Aspediotus paranerii* | *Bacteroidetes* | Silva |
| CBQZ010000001 | *Cardinium* endosymbiont cBtQ1 of *Bemisia tabaci* | *Bacteroidetes* | Silva |
| HE983995 | *Cardinium* endosymbiont cEper1 of *Encarsia pergandiella* | *Bacteroidetes* | Silva |
| AB241132 | Candidatus *Cardinium sp.* (endosymbiont of *Tetranychus urticae* red form A) | *Bacteroidetes* | Silva |
| AF350221 | Candidatus *Cardinium sp.* (endosymbiont of *Brevipalpus phoenicis*) | *Bacteroidetes* | Silva |
| AY635291 | Candidatus *Cardinium sp.* (endosymbiont of *Metaseiulus occidentalis*) | *Bacteroidetes* | Silva |
| GQ206320 | *Bacteroidetes* bacterium endosymbiont of *Sogatella furcifera* | *Bacteroidetes* | Silva |
| JX064610 | Uncultured bacterium | *Bacteroidetes* | Silva |
| AY753170 | *Bacteroidetes* endosymbiont of *Metaseiulus occidentalis* | *Bacteroidetes* | Silva |
| JN236335 | Uncultured bacterium | *Bacteroidetes* | Silva |
| JN204481 | Candidatus *Cardinium* endosymbiont of *Bemisia tabaci* | *Bacteroidetes* | Silva |
| AF319783 | *Encarsia pergandiella* asexual line endosymbiont | *Bacteroidetes* | Silva |
| AY327472 | Candidatus *Cardinium* endosymbiont of *Plagiomerus diaspidis* | *Bacteroidetes* | Silva |
| DQ854705 | Candidatus *Cardinium hertigii* | *Bacteroidetes* | Silva |
| AY928092 | Candidatus *Rhabdochlamydia crassificans* | *Chlamydiae* | GenBank |
| GAXI02038378 | Candidatus *Rhabdochlamydia sp.* (*Tetrodontophora bielanensis* (giant springtail)) | *Chlamydiae* | Silva |
| GBJM01109468 | Candidatus *Rhabdochlamydia sp.* (*Latrodectus geometricus* (brown widow)) | *Chlamydiae* | Silva |
| FPLL01005557 | Uncultured bacterium | *Chlamydiae* | Silva |
| FPLS01027258 | Uncultured bacterium | *Chlamydiae* | Silva |
| LNES01000006 | *Chlamydiae* bacterium Ga0074140 | *Chlamydiae* | Silva |
| MGME01000044 | *Chlamydiae* bacterium RIFCSPLOWO2_12_FULL_49_12 | *Chlamydiae* | Silva |
| JQ405263 | *Methanococcus maripaludis* | *Euryarchaeota* | Silva |
| KC139247 | *Methanococcus maripaludis* | *Euryarchaeota* | Silva |
| JQ178332 | *Bacillus cereus* | *Firmicutes* | Silva |
| MJEF01000067 | *Enterococcus casseliflavus* | *Firmicutes* | Silva |
| ACLZ01000007 | *Bacillus cereus* ATCC 4342 | *Firmicutes* | Silva |
| MLKF01000093 | *Bacillus cereus* | *Firmicutes* | Silva |
| JSFD01000048 | *Bacillus sp.* UMTAT18 | *Firmicutes* | Silva |
| CP002508 | *Bacillus thuringiensis serovar finitimus* YBT-020 | *Firmicutes* | Silva |
| CP001974 | *Bacillus anthracis* str. A16R | *Firmicutes* | Silva |
| CIDK01000004 | *Streptococcus pneumoniae* | *Firmicutes* | Silva |
| LSNJ01000009 | *Bacillus thuringiensis* | *Firmicutes* | Silva |
| CP010792 | *Bacillus anthracis* | *Firmicutes* | Silva |
| AJAM01000009 | *Enterococcus flavescens* ATCC 49996 | *Firmicutes* | Silva |
| BABC01000359 | Human gut metagenome | *Firmicutes* | Silva |
| ASVX01000002 | *Enterococcus flavescens* ATCC 49996 | *Firmicutes* | Silva |
| AKCC01000001 | *Enterococcus casseliflavus* EC20 | *Firmicutes* | Silva |
| AB746400 | *Wolbachia* endosymbiont of *Curculio morimotoi* | *Proteobacteria* | Silva |
| CP012420 | *Rickettsia amblyommatis* | *Proteobacteria* | Silva |
| CP018913 | *Rickettsia rickettsii* | *Proteobacteria* | Silva |
| KT175911 | *Buchnera aphidicola (Aphis gossypii)* | *Proteobacteria* | Silva |
| KT175935 | *Buchnera aphidicola (Aphis fabae fabae)* | *Proteobacteria* | Silva |
| KT175992 | *Serratia symbiotica* | *Proteobacteria* | Silva |
| LRUH01000048 | *Wolbachia* endosymbiont of *Laodelphax striatella* | *Proteobacteria* | Silva |
| MJMG01000007 | *Wolbachia pipientis* | *Proteobacteria* | Silva |
| CP001391 | *Wolbachia sp.* wRi | *Proteobacteria* | Silva |
| JX182381 | *Wolbachia pipientis* | *Proteobacteria* | Silva |
| MUJL01000115 | *Wolbachia pipientis* wUni | *Proteobacteria* | Silva |
| KX363667 | uncultured *Rickettsiaceae* bacterium | *Proteobacteria* | Silva |
| CP003884 | *Wolbachia* endosymbiont of *Drosophila simulans* wHa | *Proteobacteria* | Silva |
| DQ068812 | Uncultured bacterium | *Proteobacteria* | Silva |
| DQ068823 | Uncultured bacterium | *Proteobacteria* | Silva |
| DQ068856 | Uncultured bacterium | *Proteobacteria* | Silva |
| DQ068874 | Uncultured bacterium | *Proteobacteria* | Silva |
| GAXQ01012145 | *Wolbachia sp.* (*Pseudomasaris vespoides*) | *Proteobacteria* | Silva |
| HADL01018443 | *Wolbachia sp.* (*Calligrapha confluens*) | *Proteobacteria* | Silva |
| AP013028 | *Wolbachia* endosymbiont of *Cimex lectularius* | *Proteobacteria* | Silva |
| GCET01093322 | *Wolbachia sp.* (*Coptotermes gestroi*) | *Proteobacteria* | Silva |
| JP083175 | *Wolbachia sp.* (*Mengenilla moldrzyki*) | *Proteobacteria* | Silva |
| CP003883 | *Wolbachia* endosymbiont of *Drosophila simulans* wNo | *Proteobacteria* | Silva |
| JX669530 | Uncultured bacterium | *Proteobacteria* | Silva |
| KX155506 | *Wolbachia pipientis* | *Proteobacteria* | Silva |
| U23709 | *Wolbachia pipientis* | *Proteobacteria* | Silva |
| JMCM01000083 | *Wolbachia sp.* (*Dactylopius coccus*) | *Proteobacteria* | Silva |
| GU451192 | Uncultured bacterium | *Proteobacteria* | Silva |
| FJ774974 | *Proteobacterium* symbiont of *Nilaparvata lugens* | *Proteobacteria* | Silva |
| JHBY01090087 | *Wolbachia sp.* (*Gerris buenoi*) | *Proteobacteria* | Silva |
| LYUU01002088 | *Wolbachia sp.* ( *Armadillidium vulgare* (common pillbug)) | *Proteobacteria* | Silva |
| FPLL01008315 | metagenome | *Proteobacteria* | Silva |
| KF059257 | *Wolbachia* symbiont of *Radopholus similis* | *Proteobacteria* | Silva |
| AE006914 | *Rickettsia conorii* str. Malish 7 | *Proteobacteria* | Silva |
| CP000766 | *Rickettsia rickettsii* str. Iowa | *Proteobacteria* | Silva |
| CP003318 | *Rickettsia rickettsii* str. Hauke | *Proteobacteria* | Silva |
| AP011533 | *Rickettsia japonica* YH | *Proteobacteria* | Silva |
| LANR01000001 | *Rickettsia amblyommatis* str. Ac/Pa | *Proteobacteria* | Silva |
| CP000683 | *Rickettsia massiliae* MTU5 | *Proteobacteria* | Silva |
| CP001158 | *Buchnera aphidicola* str. Tuc7 (*Acyrthosiphon pisum*) | *Proteobacteria* | Silva |
| CP002300 | *Buchnera aphidicola* str. LL01 (*Acyrthosiphon pisum*) | *Proteobacteria* | Silva |
| CP002648 | *Buchnera aphidicola* str. Ua (*Uroleucon ambrosiae*) | *Proteobacteria* | Silva |
| CP002697 | *Buchnera aphidicola* str. USDA (*Myzus persicae*) | *Proteobacteria* | Silva |
| CP009253 | *Buchnera aphidicola* (*Aphis glycines*) | *Proteobacteria* | Silva |
| L18927 | *Buchnera aphidicola* | *Proteobacteria* | Silva |
| MKXH02000203 | *Buchnera sp.* (*Puccinia striiformi*s f. sp. Tritici) | *Proteobacteria* | Silva |
| CP013259 | *Buchnera aphidicola* (*Diuraphis noxia*) | *Proteobacteria* | Silva |
| AB642601 | Candidatus *Phytoplasma cynodontis* | *Tenericutes* | Silva |
| AF036954 | *Entomoplasma freundtii* | *Tenericutes* | Silva |
| AP014584 | *Ureaplasma parvum serovar 3* | *Tenericutes* | Silva |
| AY189305 | *Spiroplasma leptinotarsae* | *Tenericutes* | Silva |
| AY197657 | Candidatus *Phytoplasma ulmi* | *Tenericutes* | Silva |
| CP006682 | *Spiroplasma apis* B31 | *Tenericutes* | Silva |
| DQ073333 | Candidatus *Phytoplasma sp.* Guerrero LY-P2 | *Tenericutes* | Silva |
| DQ860101 | *Spiroplasma platyhelix* | *Tenericutes* | Silva |
| GU952102 | Candidatus *Phytoplasma sp.* (Coconut lethal yellowing phytoplasma) | *Tenericutes* | Silva |
| HQ613873 | Candidatus *Phytoplasma sp.* (Coconut lethal yellowing phytoplasma) | *Tenericutes* | Silva |
| JFDP01000004 | *Ureaplasma diversum* NCTC 246 | *Tenericutes* | Silva |
| JTLV01000003 | *Spiroplasma poulsonii* | *Tenericutes* | Silva |
| JX445141 | Candidatus *Phytoplasma sp.(*Bidzard witches'-broom association phytoplasma) | *Tenericutes* | Silva |
| JX843735 | *Entomoplasma ellychniae* | *Tenericutes* | Silva |
| KF801674 | Candidatus *Phytoplasma pini* | *Tenericutes* | Silva |
| KF908792 | Candidatus *Phytoplasma sp.* (Sugarcane grassy shoot phytoplasma) | *Tenericutes* | Silva |
| X80117 | Candidatus *Phytoplasma sp.* | *Tenericutes* | Silva |
| X92869 | Candidatus *Phytoplasma sp.* | *Tenericutes* | Silva |
| Y13912 | Candidatus *Phytoplasma sp.* (Cape St. Paul wilt phytoplasma) | *Tenericutes* | Silva |
| EF661582 | Candidatus *Phytoplasma sp.* (Jujube witches'-broom phytoplasma) | *Tenericutes* | Silva |
| EF186819 | Candidatus *Phytoplasma sp.* (Catharanthus phyllody phytoplasma) | *Tenericutes* | Silva |
| AM422018 | Candidatus *Phytoplasma australiense* | *Tenericutes* | Silva |
| FJ943262 | Candidatus *Phytoplasma australiense* | *Tenericutes* | Silva |
| AP006628 | Candidatus *Phytoplasma sp.* (Onion yellows phytoplasma OY-M) | *Tenericutes* | Silva |
| KU850940 | Candidatus *Phytoplasma meliae* | *Tenericutes* | Silva |
| JQ740642 | Candidatus *Phytoplasma sp.* ( 'Solanum tuberosum' phytoplasma) | *Tenericutes* | Silva |
| AJ964959 | Candidatus *Phytoplasma pyri* | *Tenericutes* | Silva |
| KX077200 | Candidatus *Phytoplasma pyri* | *Tenericutes* | Silva |
| CU469464 | Candidatus *Phytoplasma mali* | *Tenericutes* | Silva |
| KT906154 | Candidatus *Phytoplasma mali* | *Tenericutes* | Silva |
| HZ188022 | Candidatus *Phytoplasma mali* | *Tenericutes* | Silva |
| JF730310 | Candidatus *Phytoplasma prunorum* | *Tenericutes* | Silva |
| FJ432664 | Candidatus *Phytoplasma tamaricis* | *Tenericutes* | Silva |
| JQ868449 | Candidatus *Phytoplasma sp.* ('Rhamnus cathartica' stunt phytoplasma) | *Tenericutes* | Silva |
| AJ245996 | *Spiroplasma sp.* | *Tenericutes* | Silva |
| AY837731 | Uncultured *Spiroplasma sp.* | *Tenericutes* | Silva |
| FAUA01011021 | *Spiroplasma sp.* (*Heliconius timareta timareta*) | *Tenericutes* | Silva |
| KC424775 | Uncultured bacterium | *Tenericutes* | Silva |
| JN100091 | *Spiroplasma* endosymbiont of *Curculio elephas* | *Tenericutes* | Silva |
| GU993266 | *Spiroplasma platyhelix* | *Tenericutes* | Silva |
| LA837735 | *Spiroplasma sp.* (*Monomorium pharaonis* (*pharaoh ant*)) | *Tenericutes* | Silva |
| JQ768460 | *Spiroplasma sp.* crk | *Tenericutes* | Silva |
| MVYV01000081 | *Entomoplasmatales bacterium* EntAcro10 | *Tenericutes* | Silva |
| GU129148 | Uncultured *Spiroplasma sp.* | *Tenericutes* | Silva |
| JFJR01010758 | *Spiroplasma sp.* (*Glossina fuscipes fuscipes*) | *Tenericutes* | Silva |
| DQ186642 | *Spiroplasma sp.* N525 | *Tenericutes* | Silva |
| DQ319068 | *Spiroplasma kunkelii* CR2-3x | *Tenericutes* | Silva |
| X63781 | *Spiroplasma citri* | *Tenericutes* | Silva |
| GAUV02027216 | *Spiroplasma sp.* (*Acanthosoma haemorrhoidale*) | *Tenericutes* | Silva |
| AY325304 | *Spiroplasma melliferum* | *Tenericutes* | Silva |
| EF491665 | *Spiroplasma sp.* BARC 2649 | *Tenericutes* | Silva |
| CP005077 | *Spiroplasma chrysopicola* DF-1 | *Tenericutes* | Silva |
| AY189127 | *Spiroplasma chrysopicola* | *Tenericutes* | Silva |
| FAQS01006323 | *Spiroplasma sp.* (*Heliconius cydno chioneus*) | *Tenericutes* | Silva |
| FR733691 | *Spiroplasma diabroticae* | *Tenericutes* | Silva |
| CP012622 | *Spiroplasma cantharicola* | *Tenericutes* | Silva |
| AY189317 | *Spiroplasma sp.* W115 | *Tenericutes* | Silva |
| CP006681 | *Spiroplasma culicicola AES-1* | *Tenericutes* | Silva |
| AY189311 | *Spiroplasma velocicrescens* | *Tenericutes* | Silva |
| DQ163950 | Uncultured bacterium | *Tenericutes* | Silva |
| JMKV01000001 | *Mesoplasma syrphidae* ATCC 51578 | *Tenericutes* | Silva |
| JAGW01000001 | *Entomoplasma luminosum* ATCC 49195 | *Tenericutes* | Silva |
| AF547212 | *Entomoplasma lucivorax* | *Tenericutes* | Silva |
| JAGV01000003 | *Entomoplasma somnilux* ATCC 49194 | *Tenericutes* | Silva |
| AY157871 | *Entomoplasma somnilux* | *Tenericutes* | Silva |
| U26037 | *Mycoplasma mycoides subsp. capri* | *Tenericutes* | Silva |
| CP010267 | *Mycoplasma mycoides subsp. mycoides* | *Tenericutes* | Silva |
| CP002107 | *Mycoplasma mycoides subsp. mycoides* SC str. Gladysdale | *Tenericutes* | Silva |
| U26046 | *Mycoplasma capricolum subsp. capricolum* | *Tenericutes* | Silva |
| CP000123 | *Mycoplasma capricolum subsp. capricolum* ATCC 27343 | *Tenericutes* | Silva |
| CP001621 | Unidentified plasmid | *Tenericutes* | Silva |
| ANIV01000018 | *Mycoplasma mycoides subsp. capri* PG3 | *Tenericutes* | Silva |
| JN644764 | *Entomoplasma ellychniae* | *Tenericutes* | Silva |
| AY168929 | *Mesoplasma corruscae* | *Tenericutes* | Silva |
| AF042194 | *Mycoplasma monodon* | *Tenericutes* | Silva |
| AY166704 | *Mesoplasma chauliocola* | *Tenericutes* | Silva |
| DQ514605 | *Mesoplasma coleopterae* | *Tenericutes* | Silva |
| AY187288 | *Mesoplasma tabanidae* | *Tenericutes* | Silva |
| AF300327 | *Mesoplasma florum* | *Tenericutes* | Silva |
| AF305693 | *Mesoplasma entomophilum* | *Tenericutes* | Silva |
| AY174170 | *Mesoplasma grammopterae* | *Tenericutes* | Silva |
| CP006778 | *Mesoplasma florum* W37 | *Tenericutes* | Silva |
| AE017263 | *Mesoplasma florum* L1 | *Tenericutes* | Silva |
| D78650 | *Ureaplasma diversum* | *Tenericutes* | Silva |
| ABER01000002 | *Ureaplasma parvum serovar* 14 str. ATCC 33697 | *Tenericutes* | Silva |
| CP000942 | *Ureaplasma parvum serovar* 3 str. ATCC 27815 | *Tenericutes* | Silva |
| AAZQ01000001 | *Ureaplasma parvum serovar* 6 str. ATCC 27818 | *Tenericutes* | Silva |
| ABES01000001 | *Ureaplasma parvum serovar* 1 str. ATCC 27813 | *Tenericutes* | Silva |
