## Supplementary figures and images for "Tissue- and population-level microbiome analysis of the wasp spider *Argiope bruennichi* identifies a novel dominant bacterial symbiont"

### Supplementary Figure 1

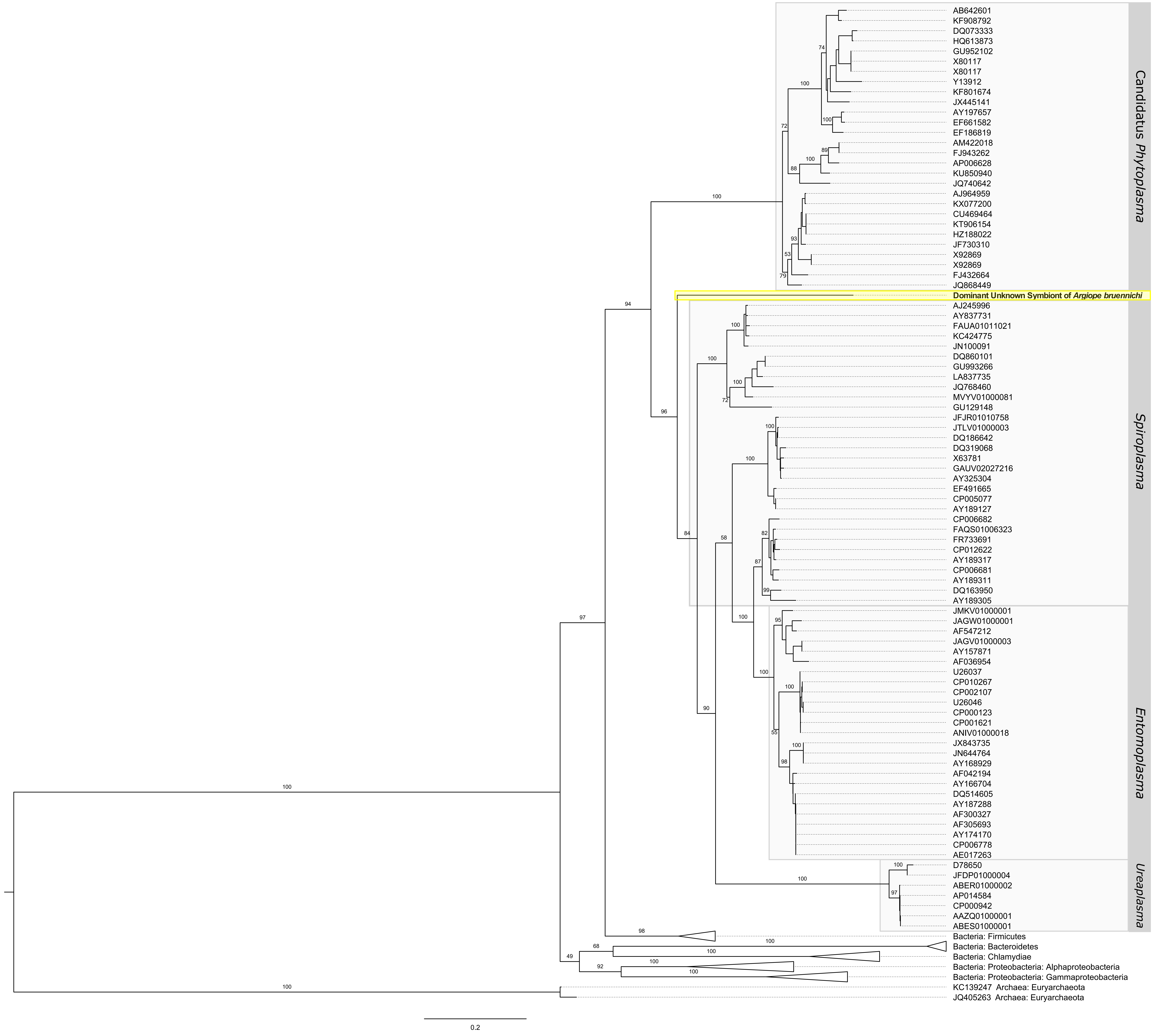
